# Multi-omic analysis of the ciliogenic transcription factor *RFX3* reveals a role in promoting activity-dependent responses via enhancing CREB binding in human neurons

**DOI:** 10.1101/2025.02.27.640588

**Authors:** Jenny Lai, Didem Demirbas, Kaitlyn Phillips, Boxun Zhao, Harrison Wallace, Megan Seferian, Tojo Nakayama, Holly Harris, Aikaterini Chatzipli, Eunjung Alice Lee, Timothy W. Yu

**Affiliations:** Division of Genetics and Genomics, Department of Pediatrics, Boston Children’s Hospital, Boston, MA, 02115, USA; The Manton Center for Orphan Disease Research, Boston Children’s Hospital, Boston, MA, 02115, USA; Broad Institute of MIT and Harvard, Cambridge, MA, 02142, USA; Program in Neuroscience, Harvard University, Boston, MA, 02115, USA; Harvard Medical School, Boston, MA, 02115, USA; Department of Pediatrics, Baylor College of Medicine and Meyer Center for Developmental Pediatrics, Texas Children’s Hospital, Houston, Texas, 77054, USA

**Keywords:** RFX, neurodevelopment, autism, human iPSC, organoids, activity-dependence, synaptic plasticity, scRNA-seq, transcriptomics, CUT&RUN-seq

## Abstract

Heterozygous loss-of-function (LoF) variants in *RFX3,* a transcription factor known to play key roles in ciliogenesis, result in autism spectrum disorder (ASD) and neurodevelopmental delay. RFX binding motifs are also enriched upstream of genes found to be commonly dysregulated in transcriptomic analyses of brain tissue from individuals with idiopathic ASD. Still, the precise functions of *RFX3* in the human brain is unknown. Here, we studied the impact of *RFX3* deficiency using human iPSC-derived neurons and forebrain organoids. Biallelic loss of *RFX3* disrupted ciliary gene expression and delayed neuronal differentiation, while monoallelic loss of *RFX3* did not. Instead, transcriptomic and DNA binding analyses demonstrated that monoallelic *RFX3* loss disrupted synaptic target gene expression and diminished neuronal activity-dependent gene expression. RFX3 binding sites co-localized with CREB binding sites near activity-dependent genes, and *RFX3* deficiency led to decreased CREB binding and impaired induction of CREB targets in response to neuronal depolarization. This study demonstrates a novel role of the ASD-associated gene RFX3 in shaping neuronal synaptic development and plasticity.

## Introduction

Autism spectrum disorder (ASD) is a highly heritable and genetically heterogeneous condition characterized by deficits in social communication and restricted, repetitive patterns of behavior. *De novo* loss-of-function variants in over 100 genes are recognized as significant contributors to ASD risk^1^. Many of these risk genes are broadly known to have roles in gene expression regulation and/or neuronal communication^1^. Studies of epigenetic and transcriptional changes in postmortem brain tissue from individuals with ASD further point to alterations in synaptic transmission as a point of functional convergence^2–4^, yet our understanding of the mechanistic links between these observations remains incomplete. Functional neurobiological characterization of ASD risk genes in experimentally manipulable human model systems could help bridge this gap.

*RFX3* encodes a member of the *RFX* family of transcription factors that is characterized by a highly conserved winged-helix DNA binding domain that recognizes the X-box motif^5^. Haploinsufficiency of *RFX3* leads to a human neurodevelopmental disorder with features including ASD, intellectual disability (ID), attention-deficit/hyperactivity disorder (ADHD), and behavioral dysregulation^6^. However, the molecular and cellular bases for these clinical impacts on human brain development and function are not understood.

Previous studies of *RFX3* in animal models have shown that *RFX3* regulates ciliogenesis^7–9^. Mouse *Rfx3* knockouts exhibit defects in ciliary assembly and function that result in hydrocephalus, situs inversus, and corpus callosum malformations^10–13^. *Rfx3* knockout mice also fail to undergo proper differentiation of auditory hair cells as well as pancreatic beta-islet cells^14,15^. Notably, our understanding of the biological roles of *RFX3* has been based on studies of animals lacking both copies of *RFX3* (i.e. complete knockouts). Prior to the discovery of the human neurodevelopmental disorder caused by heterozygous LoF variants in *RFX3*^6^, haploinsufficient phenotypes had not been described. This raised several questions with respect to the mechanistic basis for the human *RFX3* neurodevelopmental syndrome. Do symptoms observed in individuals with heterozygous *RFX3* mutations reflect previously described molecular and cellular functions of *RFX3* in ciliogenesis (noting that recessive human ciliopathies like Bardet–Biedl or Joubert Syndrome can be associated with neurocognitive impairments)? Alternatively, might the neurobehavioral impacts of *RFX3* haploinsufficiency reflect distinct biological roles or downstream targets?

To begin to address these questions, we used human iPSC-derived neuronal cultures and organoids to investigate the impacts of altering *RFX3* gene dosage on human neuronal development and function. We show that in addition to its known role as a transcriptional activator of ciliary genes, RFX3 binds cis-regulatory elements of synaptic genes. We show that RFX3 binding regulates the expression of these brain-expressed targets in a gene dosage-sensitive manner, and that single copy loss of *RFX3* (i.e., heterozygous LoF mutation) results in altered synapse formation and disrupted patterns of spontaneous neuronal activity. We further demonstrate that RFX3 facilitates CREB binding at the promoters of activity-dependent genes and is important for their induction in response to neuronal depolarization. This study provides novel insights into regulatory roles of human *RFX3* in neuronal development and synaptic plasticity and implicates alterations in tuning of activity dependent responses in human neurodevelopmental disorders.

## Results

### *RFX3* regulates neurogenesis in human forebrain organoids

*RFX3* gene expression is enriched in human embryonic brain, especially in progenitors and maturing excitatory neurons^6^. To begin to explore possible roles for *RFX3* in human brain development, we generated dorsal forebrain organoids^16^ from isogenic human iPSCs with heterozygous (HET) and homozygous (KO) loss-of-function (LoF) variants in *RFX3* CRISPR-Cas9 engineered into a control (WT) iPSC background (Figure S1A). We generated four iPSC lines bearing frameshift variants into exon 5, upstream of the DNA binding domain: HET clone 1, HET clone 2, KO clone 1, and KO clone 2, bearing heterozygous NM_001282116.2:c.313delT, heterozygous c.313insN, homozygous c.335insA, and homozygous c.335insN variants, respectively. Western blot analysis demonstrated ∼50% reduced RFX3 protein levels in HET iPSC lines, and no detectable protein in KO iPSC lines, consistent with these variants being null alleles (Figure S1B).

We cultured forebrain organoids bearing 0, 1, or 2 copies of functional *RFX3* (KO, HET, or WT). Organoids were harvested at days 45 and 90 and subject to single-cell RNA-sequencing (scRNAseq) to examine neurodevelopmental trajectories and cell type specific transcriptional changes (Figure 1A). 114,962 cells were profiled from a total of 34 control and mutant forebrain organoids across the two time points (Figure S1C-D). Overall transcriptional profiles at day 45 and day 90 correlated with the profiles of early-mid gestation stage (8-21 post-conception weeks) developing human brain samples from the Brainspan database^17^, coinciding with periods of progenitor proliferation and neurogenesis when *RFX3* expression is high^18^ (Figure S1E). *RFX3* was broadly expressed in progenitors and neurons in the organoids at these two timepoints (Figure S1F). Integration and annotation of cell types using an ensemble of references from human fetal brain and forebrain organoid atlases^19–21^ identified five major cell types (Figure 1B). At day 45, the organoids had developed cycling and outer radial glia (cycling_RG and oRG, *MKI67+, ASPM+, TNC+, PTPRZ1+*) and deep layer excitatory neurons (ExDL, *FOXP2+*). By day 90, intermediate progenitors and upper layer neurons (IP and ExUL, *SATB2+, CUX1+*) emerged (Figure 1B). Each organoid reproducibly generated each cell type, and the expression of top cell type marker genes was highly similar across individual control organoids (Figure 1C).

**Figure 1:**
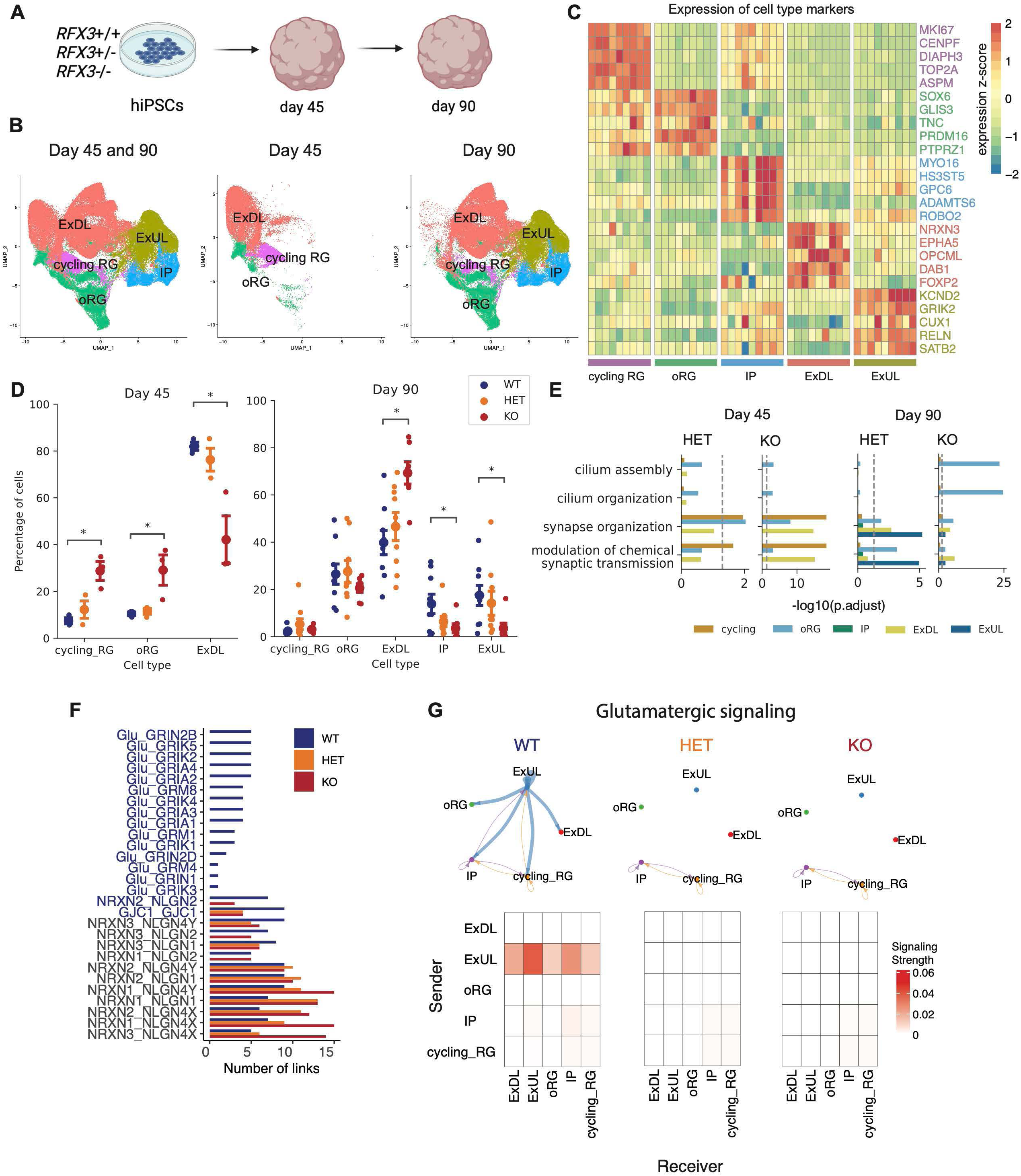
Altered glutamatergic signaling in *RFX3* deficient dorsal forebrain organoids. A. Schematic of dorsal forebrain organoid development. B. UMAP visualization of five major cell types identified in dorsal forebrain organoids over development. C. Top five marker genes per cell type in day 90 WT organoids. Each column corresponds to an individual organoid replicate (n=9). Color bar represents z-scaled expression. D. Cell type proportions per organoid at day 45 (n=3 per genotype) and day 90 (n=9 per genotype). Data represent mean +/− SEM. *p<0.05, two-sided t-test with Bonferroni correction. E. Gene Ontology (GO) term enrichment among significantly downregulated genes in *RFX3* HET and KO cell types. Benjamini-Hochberg (BH) adjusted p-values calculated with one-sided Fisher’s exact test. Dashed line indicates significance threshold, BH-adjusted p-value=0.05. F. Ranking of ligand-receptor (L-R) pairs by number of links in WT, HET, and KO organoids. L-R pairs in blue text indicate significantly increased in WT organoids compared to HET and KO (p-value<0.05, Wilcoxon test). G. Top, circle plot showing aggregated number of interactions for the significant glutamatergic L-R pairs in WT, HET, and KO organoids. Arrow width indicates number of significant interactions. Bottom, heatmap of aggregated communication strength for the significant glutamatergic L-R pairs in WT, HET, and KO organoids.

### Biallelic loss of *RFX3*, but not *RFX3* haploinsufficiency, leads to delayed neurogenesis

We asked whether *RFX3* disruption may impact developmental cell fates by comparing cell type proportions in control versus *RFX3* deficient organoids. At day 45, *RFX3* KO organoids had an increased proportion of cycling radial glia (p=0.0073) and outer radial glia (p=0.046), accompanied by a decreased proportion of deep layer neurons (p=0.018) compared to WT organoids (Figure 1D, two-sided t-test with Bonferroni correction). At day 90, *RFX3* KO organoids exhibited an increased proportion of deep layer neurons (p=0.0010) and decreased proportion of upper layer neurons (p=0.018) (Figure 1D). This suggested a temporal delay in corticogenesis in organoids with biallelic loss of *RFX3.* On the other hand, cell type proportions in *RFX3* HET organoids were indistinguishable from isogenic controls (Figure 1D), indicating a lack of detectable impact of RFX3 haploinsufficiency on cell fate specification or timing.

### Cell type-specific transcriptional alterations by *RFX3* dosage

In addition to its developmental enrichment, *RFX3* expression is maintained in the postnatal brain into adulthood and is found at especially high levels in upper layer excitatory cortical neurons that engage in cortical-cortical connectivity and demonstrate altered gene expression and function in models of ASD^3,6,22–26^. This suggested potential important biological roles in shaping neuronal function. To explore this hypothesis, we examined genes differentially expressed within each cell type in *RFX3* deficient (HET or KO) organoids compared to WT (Figure S2-S3, Table S1). Since *RFX3* is a transcriptional activator, we focused on downregulated genes (employing a false discovery rate [FDR]-adjusted cutoff of p<0.05). Genes downregulated in *RFX3* KO organoids were enriched for cilium assembly and organization genes, specifically in oRG (Figure 1E). This effect was seen at both day 45 and day 90 but was more pronounced by day 90 (FDR-adjusted p<0.05, one-sided Fisher’s exact test). (Figure 1E). In contrast, *RFX3* HET organoids did not exhibit significant dysregulation of these ciliary processes in any cell type or timepoint (Figure 1E, Table S2). Instead, genes downregulated in *RFX3* HET organoids were enriched for synapse organization, and modulation of chemical synaptic transmission (FDR-adjusted p<0.05, one-sided Fisher’s exact test) (Figure 1E). This pattern was recapitulated across several cell types and at both developmental timepoints (Figure S2-S3, Table S2). Ligand-receptor (L-R) analysis using NeuronChat^27^ further suggested decreases in glutamatergic signaling (ligand: glutamate, receptors: *GRIN2B, GRIK5, GRIK2, GRIA4, GRIA2, GRM8, GRIK4, GRIA3, GRIA1, GRM1, GRIK1, GRIN2D, GRM4, GRIN1, GRIK3)* in *RFX3* HET (and KO) organoids compared to WT (Figure 1F, blue-text indicates p-values<0.05, Wilcoxon test). In aggregate, these altered glutamatergic L-R pairs predominantly originated from ExUL in WT organoids, while fewer and weaker interactions were present between ExUL and other cell types in *RFX3* deficient organoids (Figure 1G), corresponding with the known enrichment of *RFX3* expression in this cortical layer

### Synaptic genes are sensitive to *RFX3* dosage

To explore these findings in more depth, we used *Ngn2* induction to generate *RFX3* WT, HET, and KO iPSC-derived neuronal cultures enriched for glutamatergic ExUL cell types^28^. *RFX3* WT neuronal cultures demonstrated robust RFX3 expression assessed by Western blot at day 14, while levels were decreased in HET and KO neurons as expected (Figure S4A-B). We profiled gene expression patterns of *RFX3* WT, HET, and KO neurons at day 14 in culture to identify differentially expressed genes (FDR-adjusted p<0.05). (Figure 2A, Table S3). Functional enrichment analysis revealed that, similar to dorsal forebrain organoids, genes downregulated in *RFX3* KO neurons were enriched for ciliary processes (cilium assembly, cilium organization) (FDR-adjusted p<0.05, one-sided Fisher’s exact test), whereas genes downregulated in *RFX3* HET neurons were not (Figure 2B, Table S4-S5). Overall expression of a ciliary gene set (CiliaCarta, n=956) was also on average significantly reduced in *RFX3* KO neurons (p<0.05, one-sided permutation test) (Figure 2C). The functional significance of these findings was corroborated by immunostaining for ARL13B, a ciliary GTPase mutated in Joubert syndrome (70% reduced protein levels in RFX3 KO levels, p<0.05, one-way t-test with Bonferroni correction, Figure S4C-F). By contrast, no changes in any of these measures were detected in *RFX3* HET neurons (Figure 2B, Figure 2C, Figure S4C-F).

**Figure 2:**
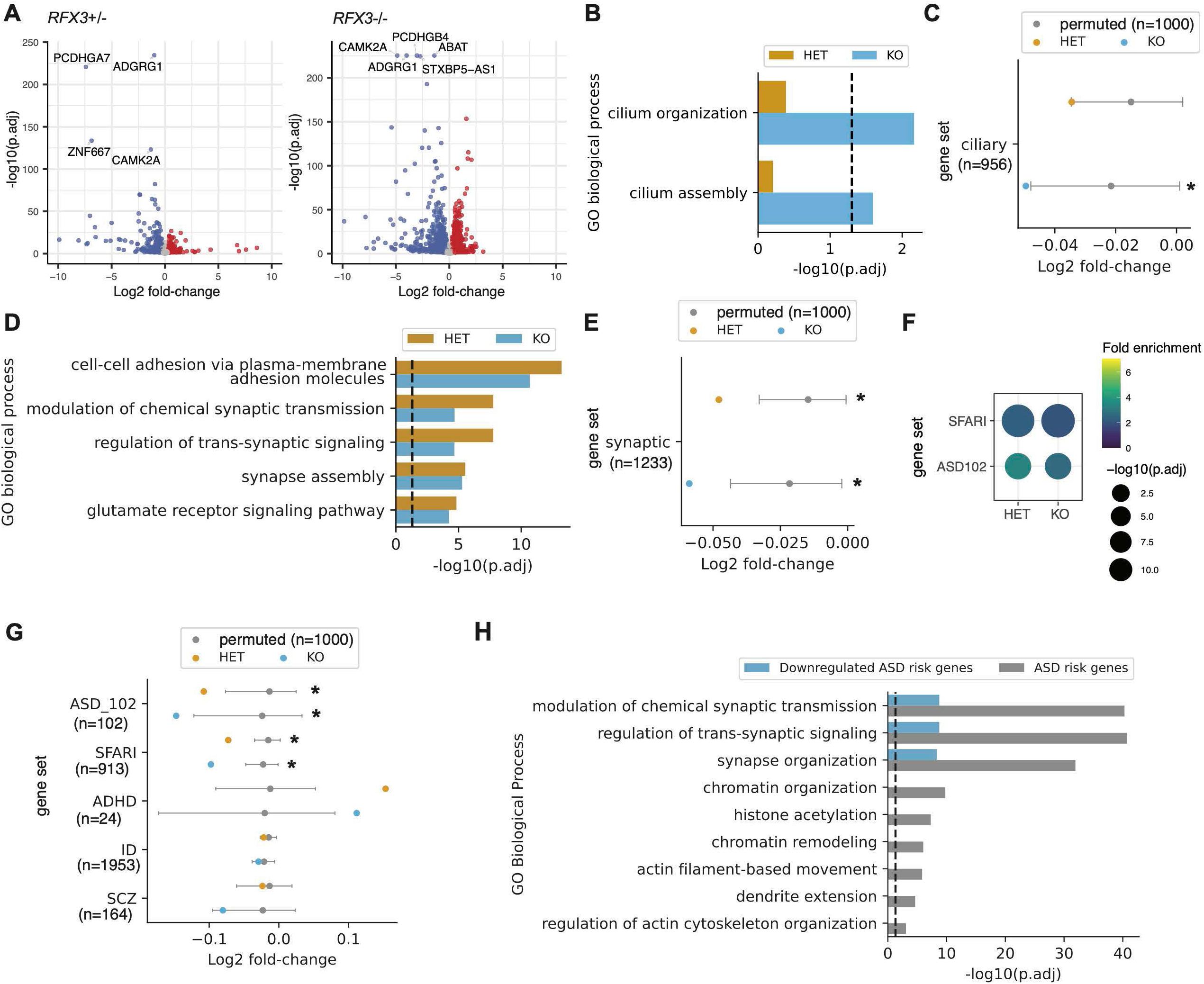
Synaptic gene expression is sensitive to RFX3 dosage. A. Volcano plot showing differentially expressed genes (DEGs) between *RFX3* HET and WT neurons (left) and *RFX3* KO and WT neurons (right) at day 14 in culture. Genes significantly downregulated are in blue (log2FC<-0.25, FDR<0.05). Genes significantly upregulated are in red (log2FC>0.25, FDR<0.05). The top DEGs ranked by significance are labeled. B. Gene Ontology (GO) term enrichment analysis of significantly downregulated genes in *RFX3* HET and KO. Ciliary related GO terms shown. Benjamini-Hochberg (BH) adjusted p-values calculated with one-sided Fisher’s exact test. Dashed line indicates significance threshold, BH-adjusted p-value=0.05. C. Average observed log2 fold-change of ciliary gene set (n=956 CiliaCarta) in *RFX3* HET and KO neurons vs WT neurons compared to permuted distribution for randomly sampled gene sets of equal size (n permutations=1,000, 90 percentile interval). *left-tailed p <0.05, permutation test. D. Gene Ontology (GO) term enrichment analysis of significantly downregulated genes in *RFX3* HET and KO. Top enriched GO terms related to neurodevelopment shown. Benjamini-Hochberg (BH) adjusted p-values calculated with one-sided Fisher’s exact test. Dashed line indicates significance threshold, BH-adjusted p-value=0.05. E. Average observed log2 fold-change of synaptic gene set (n=1233 SynGO) in *RFX3* HET and KO neurons compared to permuted distribution for randomly sampled gene sets of equal size (n permutations=1,000, 90 percentile interval). *left-tailed p <0.05, permutation test. F. Overrepresentation analysis of ASD risk genes among *RFX3* HET and KO downregulated genes. Dot size indicates significance of enrichment. Color indicates fold enrichment (fraction of overlapping genes compared to probability of obtaining overlap by chance). G. Average observed log2 fold-change of neuropsychiatric risk gene sets in *RFX3* HET and KO neurons vs WT neurons compared to permuted distribution for randomly sampled gene sets of equal sizes (n permutations=1,000, 90 percentile interval). *left-tailed p <0.05, permutation test. ID, intellectual disability. SCZ, schizophrenia. ADHD, attention-deficit hyperactivity disorder. H. Gene Ontology (GO) term enrichment analysis of all ASD risk genes (ASD102 and SFARI union set, n=938) and those significantly downregulated in *RFX3* HET and KO neurons. Dashed line indicates significance threshold FDR 0.05.

Amongst genes downregulated in *RFX3* HET neurons, the top enriched pathways were related to neurodevelopment and synaptic function, including forebrain development, modulation of chemical synaptic transmission, synapse assembly, and synapse organization (FDR-adjusted p<0.05, one-sided Fisher’s exact test) (Figure 2D, Table S4-S5). Synaptic genes as a set (SynGO^29^, n=1,233) had significantly reduced expression in *RFX3* HET neurons as well (p<0.05, one-sided permutation test) (Figure 2E). Genes downregulated in *RFX3* HET neurons were also enriched for ASD risk genes, defined by SFARI (n=913) and a recent large exome sequencing study (ASD102^1^, n=102) (FDR-adjusted p<0.05, one-sided Fisher’s exact test) (Figure 2F). We also analyzed the overall expression of ASD risk gene sets as well as gene sets for known genetic causes of intellectual disability (ID) (ORPHA:183757, n=1,953), ADHD (n=24)^30–32^, and schizophrenia (SCZ, n=164)^33^. ASD risk genes (ASD102 and SFARI sets) were significantly downregulated in *RFX3* HET neurons (p<0.05, one-sided permutation test), while ADHD, ID, and SCZ risk gene sets were not (Figure 2G). Supporting the specificity of these findings, all ASD risk genes (ASD102 and SFARI union set, n=938 genes) were enriched for synaptic, cytoskeletal, and chromatin processes, but those with decreased expression in *RFX3* HET neurons were only enriched for synaptic processes (FDR-adjusted p<0.05, one-sided Fisher’s exact test), suggesting that *RFX3* regulates a subset of ASD risk genes primarily involved in neuronal communication (Figure 2H). These results corroborated the altered glutamatergic synaptic signaling in *RFX3* deficiency observed in our organoid analysis.

### *RFX3* exhibits dosage sensitive binding near synaptic genes

In principle, downregulation of synaptic genes in *RFX3* deficient neurons could reflect direct effects of decreased RFX3 binding to the cis-regulatory elements of synaptic genes, or indirect effects of decreased RFX3 binding to genes responsible for specifying neuronal cell development (e.g. differentiation/maturation/fate). Morphological analysis did not reveal detectable differences in total neurite length (marked by TUJ1) in *RFX3*+/− neurons (Figure S5A). Furthermore, RNA deconvolution analysis estimating the maturity of progenitors and neurons in *RFX3* WT, HET, and KO iPSC-derived neuronal cultures^34^ did not show differences in fractions of progenitors and neurons between genotypes over multiple time points (Figure S5B).

To directly identify RFX3 targets in human neurons, we profiled RFX3 binding genome-wide using CUT&RUN-sequencing. This yielded 4,240 RFX3-specific binding peaks (peaks present in WT neurons and not present in KO neurons, Figure 3A, Table S6, n=4,303 peaks in WT, n=287 peaks in KO). The *RFX3* motif was the most enriched among these binding regions and was present in 86% of peaks (E-value 3.70e-922, Figure 3B). In addition, binding motifs for SIX1 (AACCTGA) (E-value 5.11e-7) and CREB (TGACGTCA) (E-value 2.02e-6) were identified as the top enriched co-occurring motifs (Figure 3C). SIX1 and RFX3 have been shown to cooperatively interact to regulate gene expression programs in auditory sensory epithelium^35^, and RFX5 (a close relative of RFX3 with similar binding specificity) and CREB cooperatively bind the X2 box in MHC class II promoters^36^, suggesting that RFX3 may cooperate with these transcription factors in human neurons as well. Genes with RFX3-specific binding peaks were enriched for ciliary related and synaptic related functions (Figure 3D). Consistent with RFX3 acting as a direct transcriptional activator, RFX3 target genes exhibited significantly decreased expression in *RFX3* deficient (HET or KO) neurons in a gene dosage-dependent fashion (Bonferroni-adjusted p<0.05, t-test) (Figure 3E). Promoter peaks (within 3,000 bp of the adjacent gene transcription start site) were enriched for both ciliary processes and synaptic processes, while distal peaks (>3,000 bp) were enriched for synaptic processes (FDR-adjusted p<0.05, one-sided Fisher’s exact test) (Figure 3D).

**Figure 3:**
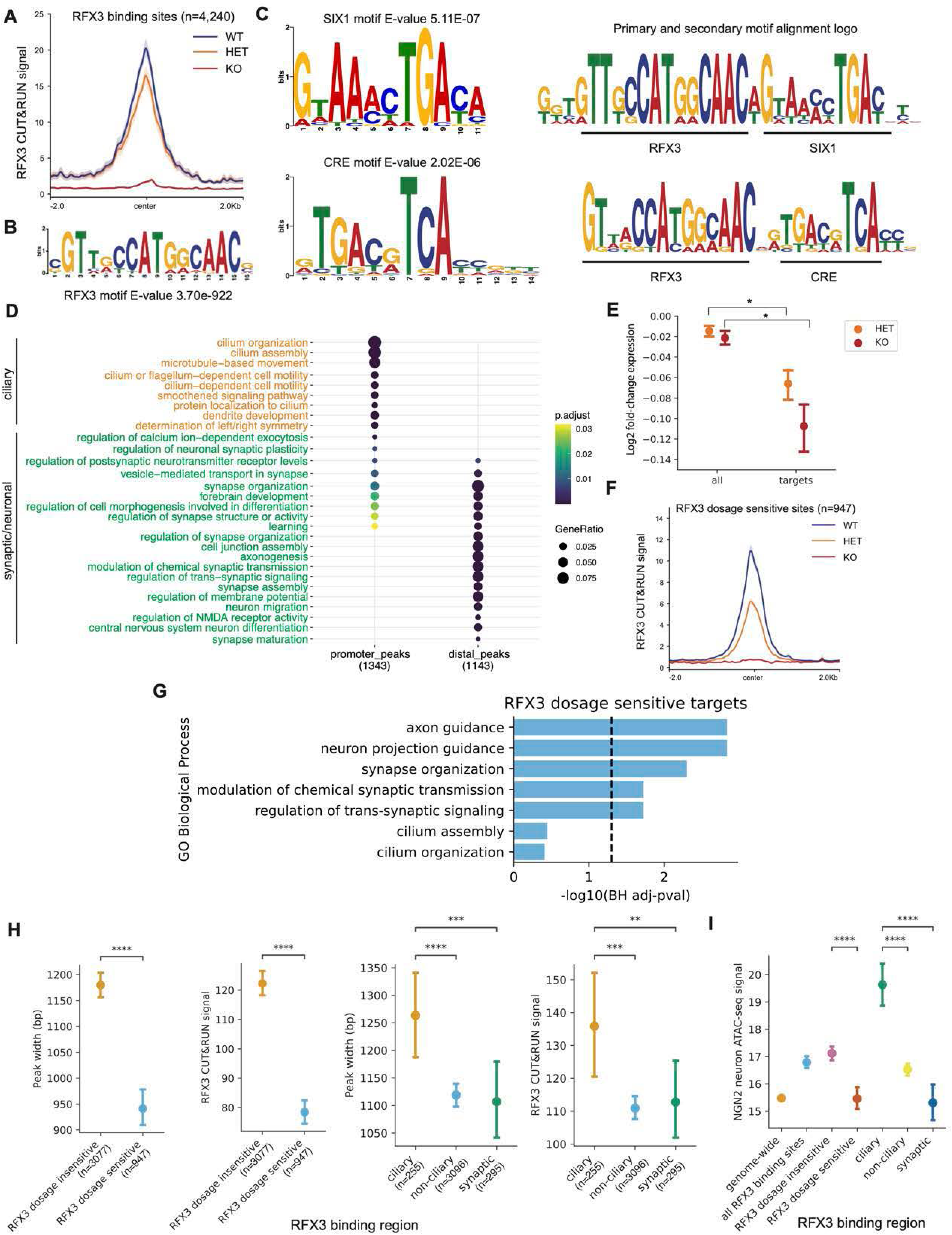
*RFX3* exhibits dosage sensitive binding near synaptic genes. A. Aggregate plot of RFX3 CUT&RUN reads within RFX3 binding peaks called in WT neurons (n=4,240 peaks). Trace shows mean +/− SEM of n=7 WT replicates, n=6 HET replicates, n=6 KO replicates. B. RFX3 binding motif enriched within RFX3 binding peaks. C. SIX1 binding motif enrichment near RFX3 binding motifs in RFX3 binding peaks. Primary (RFX3) and secondary (SIX1) motif alignment. CRE binding motif enrichment near RFX3 binding motifs in RFX3 binding peaks. Primary (RFX3) and secondary (CRE) motif alignment. D. Gene Ontology (GO) enrichment analysis of genes with an RFX3 peak in the promoter (promoter_peaks) or distal regions (distal_peaks). E. Log2 fold-changes in expression levels of all genes or RFX3 target genes in RFX3 HET or KO neurons compared to WT. *p<0.05, t-test with Bonferroni correction. F. Aggregate plot of RFX3 CUT&RUN reads within RFX3 binding peaks significantly decreased in *RFX3* HET neurons (n=947 peaks, FDR<0.2). Trace shows mean +/− SEM of n=7 WT replicates, n=6 HET replicates, n=6 KO replicates. G. Gene Ontology (GO) enrichment analysis of *RFX3* target genes with significantly decreased peak and expression in *RFX3* HET neurons. Dashed line indicates significance threshold FDR 0.05. H. Peak width and RFX3 CUT&RUN signal among *RFX3* dosage sensitive, *RFX3* dosage insensitive, ciliary, non-ciliary, and synaptic regions. *p<0.05, **p<0.01, ***p<0.005, ****p<0.0001, t-test. I. ATAC-seq signal in *Ngn2* neurons in different sets of RFX3 binding regions. Data represent mean +/− 95% CI. *p<0.05, **p<0.01, ***p<0.005, ****p<0.0001, one-way t-test with Bonferroni correction.

Next we characterized how RFX3 binding patterns change in the heterozygous mutant state. We found that 947/4,240 RFX3 binding sites (22%) demonstrated significantly decreased binding (FDR-adjusted p<0.2) in *RFX3* HET neurons compared to WT (Figure 3F, Table S7); 124/947 of these (13%) exhibited decreased expression as well. These dosage sensitive RFX3 targets were enriched for axon guidance (i.e., *PLXNC1, NOG, BMPR1B*) and synapse organization (i.e., *CACNB2, SRGAP2, GRIN2B, CBLN2*) (FDR-adjusted p<0.05, one-sided Fisher’s exact test) pathways (Figure 3G). Dosage-sensitive RFX3 targets were not enriched for ciliary pathways (Figure 3G).

### *RFX3* has weaker binding near synaptic genes associated with decreased chromatin accessibility

We explored why RFX3 binding sites near synaptic targets might be especially sensitive to genetic dosage, relative to binding sites near ciliary genes. In WT neurons, *RFX3* dosage insensitive sites had significantly increased peak width and height compared to dosage sensitive sites (p<0.0001, t-test) (Figure 3H). Ciliary genes also had RFX3 peaks with increased peak width and height compared to non-ciliary genes and synaptic genes (p<0.01, t-test) (Figure 3H). This suggested that RFX3 binding sites that are sensitive to genetic dosage may simply be those with weaker RFX3 binding at baseline. To investigate this further, we also considered whether RFX3 binding strength may be influenced by additional factors. Ciliary and synaptic targets contained a similar number of RFX3 motifs (∼2 RFX3 motifs per binding region; Figure S5C), and exhibited comparable RFX3 motif scores (∼12, Figure S5D). In contrast, analysis of human neuronal ATAC-seq data^37^ showed that *RFX3* dosage sensitive sites exhibited significantly decreased chromatin accessibility compared to dosage insensitive sites (Bonferroni-adjusted p<0.0001, t-test) (Figure 3I). RFX3 binding sites near synaptic genes also exhibited decreased chromatin accessibility compared to binding sites near ciliary genes (Bonferroni-adjusted p<0.0001, t-test) (Figure 3I), implicating chromatin accessibility as a key contributing factor rendering synaptic targets especially sensitive to *RFX3* gene dosage.

### Reduced synapse number and delayed neuronal network maturation in *RFX3* haploinsufficiency

We considered whether the transcriptional deficits observed in *RFX3* deficient neurons have functional consequences on synaptic assembly or function. Excitatory synapse numbers, quantified by SYN1 and PSD-95 colocalization, were significantly decreased in day 14 *RFX3* HET neuronal cultures compared to control (p<0.05, one-way t-test with Bonferroni correction) (Figure 4A). We used multielectrode arrays (MEA) to measure spontaneous synaptic activity of *RFX3* deficient neurons compared to isogenic WT controls from one week to eight weeks in culture. MEA captures complex neuronal network activity over time, where neuronal activity progresses from random spikes, to electrode bursts, to organized patterns of network wide bursts. Synchronized firing represents strong synaptic connections and plays a role in activity-dependent organization of cortical networks^38–40^. Synchronized neural networks increase over brain development and are a hallmark of mature cortical circuits^41–43^. Overall activity, measured by mean firing rate, did not differ between *RFX3* WT and HET neurons (Figure 4B). The number of spikes per network burst and synchrony were decreased in *RFX3* HET and KO neurons from day 39 to day 62 in culture (p<0.05, one-sided t-test), indicating decreased synaptic strength and immature network activity (Figure 4C-D). *RFX3* HET and KO neurons also exhibited significantly decreased neural activity scores (NAS^46^) compared to WT from day 39 onwards (p<0.05, one-sided t-test) (Figure 4E). These results demonstrate that dysregulated synaptic gene expression in *RFX3* deficient neurons is functionally associated with altered synaptic number, decreased synaptic strength, and delayed maturation of network activity.

**Figure 4:**
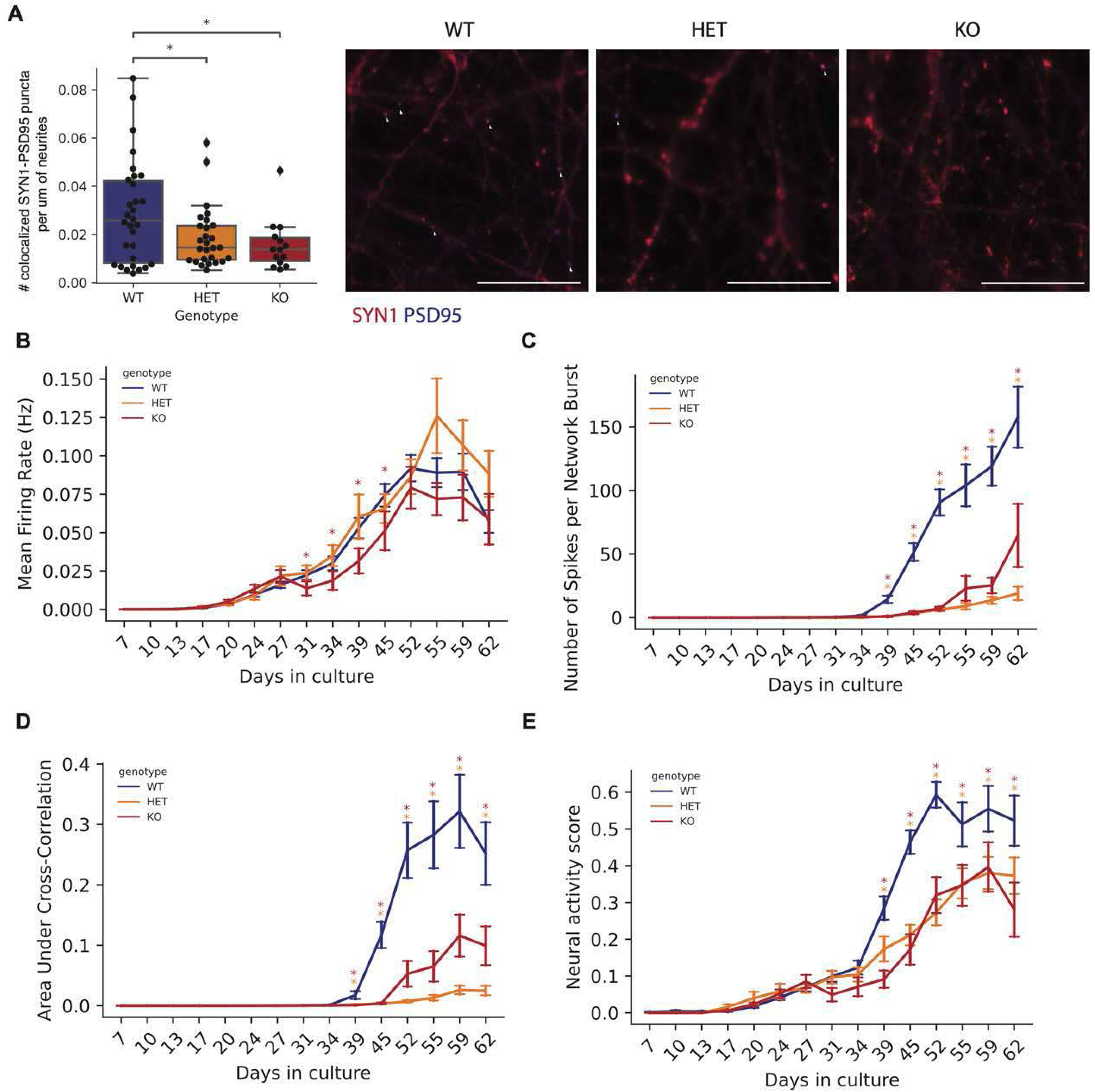
Decreased synchronized neural networks in *RFX3* deficient neurons. A. Quantification and representative images of SYN1 and PSD95 colocalized puncta on neurites in day 14 neuronal cultures. n=32 wells per genotype. White arrows mark representative colocalized puncta. Scale bar=100 um. *p<0.05, one-way t-test with Bonferroni correction. B. Mean firing rate (Hz) of neuronal cultures from day 7 to day 62. C. Average number of spikes per network burst of neuronal cultures from day 7 to day 62. D. Area under cross-correlation (measure of synchrony) of neuronal cultures from day 7 to day 62. E. Neural activity score of neuronal cultures from day 7 to day 62. (B-E) Data normalized to the number of covered electrodes per well. Data represented as mean +/− SEM. n=32 wells per genotype from two independent MEA plates. *p<0.05, one-sided t-test, red * indicates KO vs. WT, yellow * indicates HET vs. WT.

### RFX3 modulates CREB-dependent activity-dependent transcriptional responses

RFX3 binding sites were enriched for the binding motif of CREB, a key regulator of activity dependent gene expression programs^44^ (Figure 3C). Notably, we found that RFX3 bound near the immediate early genes *FOSB* and *JUNB,* known CREB target genes involved in activity-dependent signaling pathways^45^ (Figure 5A). Furthermore, we found both the *RFX3* binding motif and RFX3 binding sites to be significantly enriched in KCl inducible H3K27ac regions in *Ngn2* neurons that mark activity-dependent regulatory elements^37^ (RFX3 motif E-value 4.12e-44; FDR-adjusted p-value<0.05, two-sided Fisher’s exact test) (Figure S6A-B). These observations suggested that *RFX3* may be a direct regulator of activity-dependent gene expression programs.

**Figure 5:**
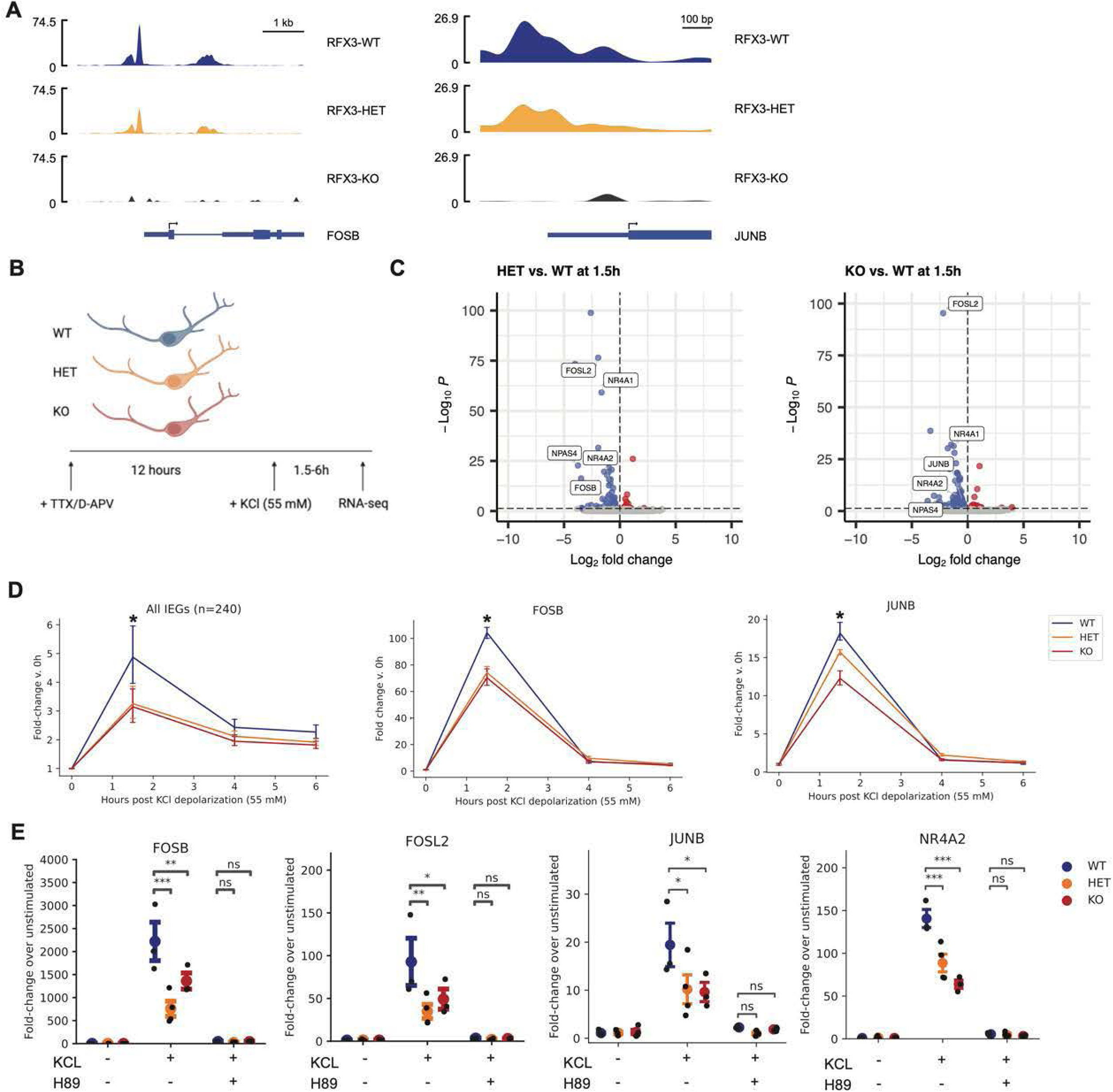
RFX3 modulates CREB-dependent activity-dependent transcriptional responses. A. RFX3 binding peaks in the promoter regions of *FOSB* and *JUNB* in *RFX3* WT, HET, and KO day 14 neurons. B. Schematic of neuronal stimulation with KCl. C. Volcano plot of differentially expressed genes in 1.5 hour KCl stimulated HET vs. WT or KO vs. WT neurons at day 14 in culture. Genes significantly downregulated are in blue and genes significantly upregulated are in red (|log2FC|>1, FDR<0.05). Select immediate early genes are labeled. D. Fold-induction of all immediate early genes, *FOSB,* and *JUNB* at 1.5, 4, and 6 hours post KCl depolarization compared to unstimulated neurons. Data represented as mean and 95% CI, *p<0.05 t-test, n=3 independent neuron differentiations per genotype. E. Fold-induction of *FOSB, FOSL2, JUNB, NR4A2* at 1.5 hours post KCl depolarization or H89 pretreatment and KCl depolarization assessed by RT-qPCR. Data represented as mean +/− SEM, n=3-4 independent wells per condition. *p<0.05, **p<0.01, ***p<0.005, ns not significant, one-way t-test with Bonferroni correction.

We used RNA-seq to profile activity-dependent gene expression responses of day 14 *RFX3* WT, HET, and KO neurons (Figure 5B). Depolarization with 55 mM of KCl results in a peak in immediate-early gene (IEG) induction after 1.5 hours and late response gene (LRG) induction after 6 hours in human iPSC-derived neuronal cultures^37,46^. Under these conditions, we identified 240 IEGs and 481 LRGs that were upregulated with KCl stimulation (fold change of at least 2, FDR-adjusted p<0.05) (Figure S6C, Table S8). IEG upregulation was significantly blunted in *RFX3* HET or KO (Figure 5C, Figure S6D). IEGs as a group showed decreased induction by ∼40% in *RFX3* deficient neurons 1.5 hours post depolarization (p<0.05, t-test) (Figure 5D). 40/240 (17%) IEGs, including AP-1 complex members *FOSB, FOSL2, JUNB,* and *JUND,* demonstrated impaired induction in both *RFX3* HET and KO neurons (FDR-adjusted p<0.05) (Figure 5D, Table S9). The degree of impaired induction was comparable to levels observed in neurons with knockdown of *CRTC1*, a CREB transcriptional coactivator^46^. *RFX3* deficient neurons also had decreased induction of LRGs (∼15%) 6 hours post depolarization (p<0.05, t-test) (Figure S6D-E).

### RFX3 promotes CREB binding at cis-regulatory elements of activity-dependent genes

We sought to explore how *RFX3* regulates induction of activity dependent genes. Depolarization did not significantly affect RNA expression levels of *RFX3* itself (Figure S6F). Furthermore, RFX3 immunostaining demonstrated a nuclear localization pattern that was unchanged by KCl stimulation (Figure S6G), suggesting that *RFX3* activity is not regulated by neuronal activity. The effects of *RFX3* on activity dependent transcription could however be blocked with treatment of H89, a protein kinase A (PKA) inhibitor that blocks CREB Ser-133 phosphorylation^47^ (Figure 5E). We also treated neurons with forskolin (FSK), a compound that activates CREB via activation of adenylate cyclase^47^. *RFX3* deficient neurons showed decreased induction of IEGs in response to FSK (Bonferroni-adjusted p<0.05, t-test) (Figure S6H).

These results suggested that RFX3 may modulate IEG induction via interactions with activated CREB. In principle, RFX3 could be required for CREB activation, or alternatively, RFX3 may act in parallel or downstream of activated CREB to modulate IEG induction. To distinguish between these two possibilities, we assessed CREB activation levels marked by phosphorylation at Ser-133 (P-CREB) in response to KCl in *RFX3* deficient compared to WT neurons. By both Western blot and immunocytochemistry, P-CREB levels at 15 min and 1.5 hours post KCl depolarization were similar between WT and *RFX3* deficient neurons (Figure S7A-D), indicating that CREB phosphorylation itself was unaffected by *RFX3* deficiency. We therefore considered whether RFX3 might act in parallel or downstream of activated CREB. We profiled CREB binding with CUT&RUN-seq in *RFX3* WT, HET, and KO unstimulated neurons. We identified 3,095 CREB peaks in WT neurons. Motif enrichment analysis revealed that the CRE motif was the most enriched among these binding regions (CRE motif E-value 5.57e-200, Figure 6A-B). In addition, the *RFX3* motif was significantly enriched and present in 10.4% of peaks (RFX3 motif E-value 4.14e-44, Figure 6B), suggesting that CREB and RFX3 might co-bind at least a subset of sites. 567 binding sites were shared between CREB and RFX3, including the promoters of AP-1 components *FOSB* and *JUNB* (Figure 6C, Table S10). On average, RFX3 co-occupied sites had 1.77-fold higher CREB CUT&RUN signal compared to sites without RFX3 (Bonferroni-adjusted p<0.001, t-test) (Figure 6D). Moreover, the average CREB peak intensity at the 567 co-bound sites was significantly decreased in *RFX3* HET neurons compared to a permuted distribution of CREB peaks in *RFX3* HET (p=0.002, permutation test, n=1000 permutations) (Figure 6E-F). These data suggest that RFX3 directly promotes CREB binding at co-bound sites. In addition, 159/567 (28%) of the CREB sites co-occupied by RFX3 were very sensitive to *RFX3* dosage, where a CREB binding peak was not called in the *RFX3* HET neurons (Table S11); these included the AP-1 complex components *FOSB* and *JUNB* (Figure 6G). Overall, these data support a mechanism whereby RFX3 and CREB are binding partners at the promoters of a subset of CREB targets. At these sites, RFX3 promotes CREB binding and *RFX3* deficiency leads to decreased CREB binding and impaired induction of *CREB* targets in response to stimulation (Figure 7).

**Figure 6:**
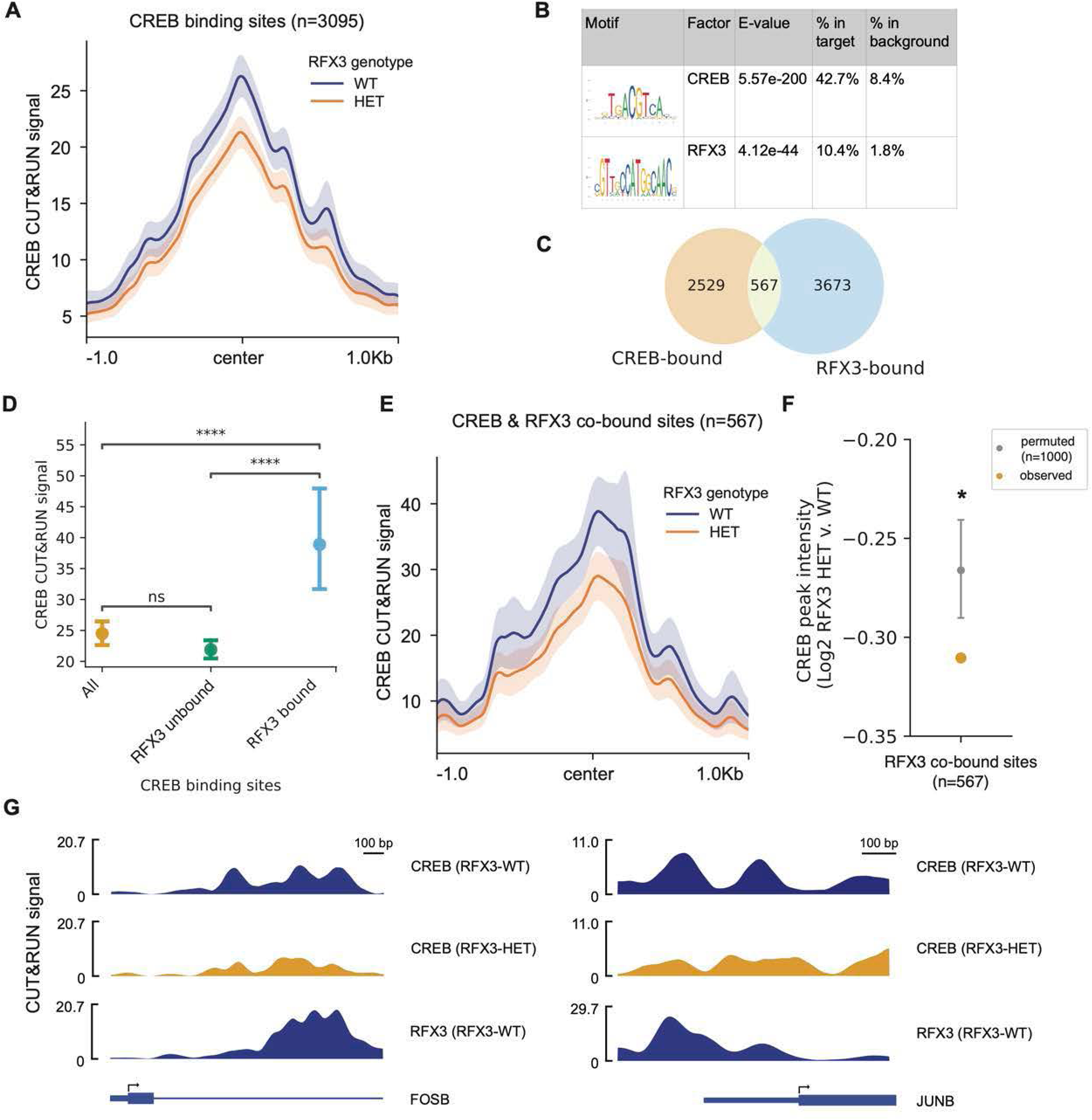
RFX3 promotes CREB binding in unstimulated neurons. A. Aggregate plot of CREB CUT&RUN reads within CREB binding peaks called in WT neurons (n=3095 peaks). Trace shows mean +/− SEM of n=2 WT replicates, n=2 *RFX3* HET replicates. B. CREB and RFX3 binding motifs enriched within CREB binding regions in WT neurons. C. Overlap between CREB-bound and RFX3-bound regions in WT neurons. D. CREB CUT&RUN signal at all, RFX3 unbound, and RFX3-bound CREB binding sites in WT neurons. ****p<0.001, ns not significant, one-way t-test with Bonferroni correction. E. Aggregate plot of CREB CUT&RUN reads within CREB and RFX3 co-bound sites (n=567 peaks). Trace shows mean +/− SEM of n=2 WT replicates, n=2 *RFX3* HET replicates. F. Average observed log2 fold-change in CREB peak intensity at regions co-bound by RFX3 in *RFX3* HET neurons compared to a permuted distribution (n permutations=1,000, 90 percentile interval). *left-tailed p <0.05, permutation test. G. CREB binding peaks near *FOSB,* and *JUNB* in *RFX3* WT and HET neurons. RFX3 binding peaks near *FOSB* and *JUNB* in *RFX3* WT neurons.

**Figure 7:**
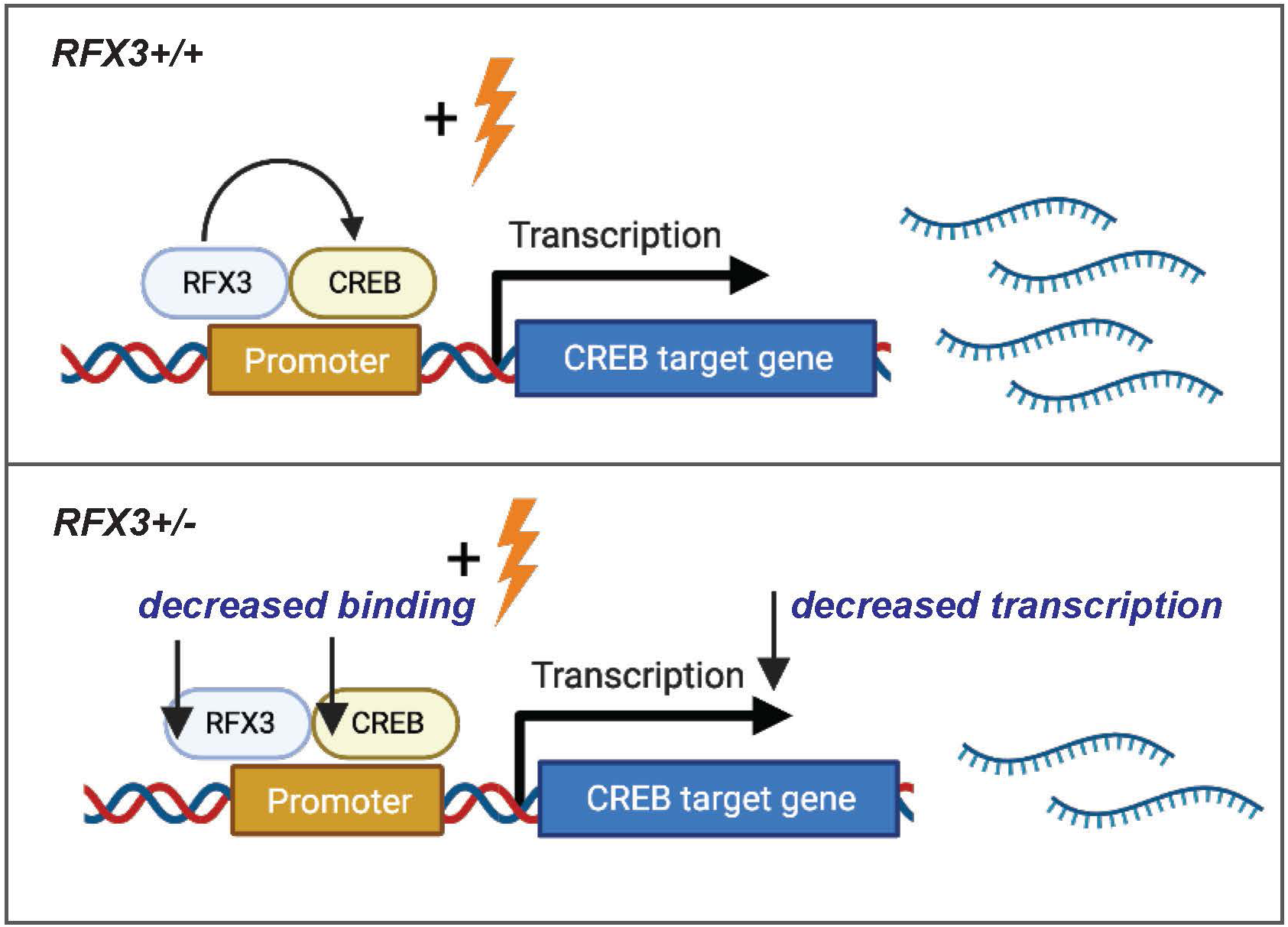
RFX3 regulates activity-dependent transcription by promoting CREB binding. Schematic of proposed mechanism where RFX3 promotes CREB binding to modulate expression of activity dependent genes.

## Discussion

*RFX3* is a brain-enriched ciliogenic transcription factor for which genetic haploinsufficiency has been associated with ASD. Here, we performed genome-wide profiling of RFX3 binding and transcriptomic alterations in *RFX3* deficiency in human neurons to provide molecular insights into the biological pathways regulated by *RFX3* and their links to ASD. Using human dorsal forebrain organoids, we found that biallelic loss of *RFX3* leads to altered neurogenesis associated with downregulation of ciliary genes. Disruption to primary cilia has been linked to impaired neurogenesis related to sonic hedgehog signaling and cell proliferation^48–51^. For example, loss of the ciliary gene *Inpp5e* (mutated in some forms of Joubert Syndrome) altered neurogenesis with overproduction of layer V neurons in mouse cortex^52^, and mutations in the gene encoding AHI1, a ciliary transition zone protein, can present with polymicrogyria characterized by abnormal cortical lamination^53,54^. In contrast, impaired neurogenesis and alterations in ciliary gene expression were not observed in *RFX3* haploinsufficiency. Instead, we show that *RFX3* haploinsufficiency leads to decreased expression of synaptic genes, including other ASD risk genes. *RFX3* deficiency was functionally associated with altered development of neuronal network activity, including decreased synchrony and number of spikes per network burst. Notably, decreased spontaneous inter-hemispheric synchrony was found in toddlers with ASD^55^. Decreased spiking activity and burst frequency has been reported in patient-derived neurons from individuals with idiopathic ASD as well as with monogenic forms of ASD^56–60^ and may correlate with reductions in functional brain connectivity and synchrony broadly reported in clinical studies of individuals with ASD^61^.

We further demonstrate that RFX3 deficiency leads to impaired induction of activity dependent genes due to reduced facilitation of CREB binding. Symptoms of neurodevelopmental disorders appear in early postnatal life, coinciding with periods of activity-dependent synapse development and refinement^62^, and dysregulation of activity dependent signaling has been previously implicated in neurodevelopmental disorders^63^. CREB-mediated activity dependent transcription is important for synapse maturation and dendritic arborization involved in learning^62^, and pathogenic genetic variation in several key components of activity dependent signaling lead to neurodevelopmental disorders, including L-VSCC (Timothy syndrome), RSK2 (Coffin-Lowry), CBP (Rubenstein-Taybi syndrome), UBE3A (Angelman), MECP2 (Rett syndrome), and BAF complex subunits that mediate AP-1 transcriptional activity^63–65^. Haploinsufficiency of *NR4A2* and *FOSL2* has recently been associated with neurodevelopmental disorders^66,67^. Therefore, impairment in activity-dependent synapse maturation may be common to genetically heterogeneous neurodevelopmental disorders.

Here, we found that RFX3 and CREB co-occupy the promoters of a subset of activity dependent response genes, and that RFX3 promotes CREB binding at these sites, suggesting that *RFX3* is a direct component of activity-dependent transcriptional complexes. RFX3 and CREB also showed evidence of protein-protein interaction in a large-scale mammalian two hybrid screen conducted in human cells^68^. RFX3 co-occupation of a subset of CREB binding sites may reflect cell-type specific modulation of activity-dependent gene programs^65,69^.

The impact of RFX3 deficiency on activity-dependent gene expression may contribute to ASD phenotypes in patients with *RFX3* deficiency and might have mechanistic implications for idiopathic ASD as well. ASD epigenetic signatures marked by increased histone acetylation (H3K27ac) were most enriched for the RFX motif and AP-1 motif^2^. In addition, CREB and RFX family motifs were enriched in activity-inducible promoters associated with elevated ASD-heritability^46^, raising the possibility that RFX may be a portion of a common downstream activity-dependent pathway that is dysregulated in idiopathic ASD.

## Acknowledgements

We thank Mandovi Chatterjee, Ph.D. from the Harvard Single Cell Core for her assistance with scRNA-seq library preparation, Lee Barrett, Ph.D., Elizabeth Buttermore, Ph.D., and Nina Makhortova, Ph.D. from the Boston Children’s Human Neuron Core for their assistance with high-content imaging and multi-electrode array assays, the Harvard Biopolymer Facility for bulk RNA-sequencing library preparation and sequencing, Christopher A. Walsh, M.D., Ph.D., Michael E. Greenberg, Ph.D., Lisa V. Goodrich, Ph.D., GiHun Choi, Ph.D, and members of the Yu and Lee labs for helpful scientific discussions. Illustrations were created with BioRender.com.

This work was supported by the National Institute of Health grant R01MH113761 and the G. Harold & Leila Y. Mathers Foundation. J.L is supported by award Number T32GM007753 and T32GM144273 from the National Institute of General Medical Sciences, and the Ruth L. Kirschstein NRSA F30 Fellowship (1F30MH128995). B.Z. was supported by the Manton Center Pilot Project Award and Rare Disease Research Fellowship. The content is solely the responsibility of the authors and does not necessarily represent the official views of the National Institute of General Medical Sciences or the National Institutes of Health.

## Author Contributions

Conceptualization, T.W.Y, E.A.L, and J.L; Methodology, T.W.Y, E.A.L, J.L, D.D; Investigation, J.L, D.D, K.P, H.W, B.Z, M.S, T.N, H.H; Formal Analysis, J.L, B.Z, A.C; Validation, K.P, H.W; Supervision, T.W.Y, E.A.L; Writing – Original Draft, J.L; Writing – Review & Editing, All.

## Inclusion and diversity

We support inclusive, diverse, and equitable conduct of research.

## Supplemental Figure Legends

**Figure S1:**
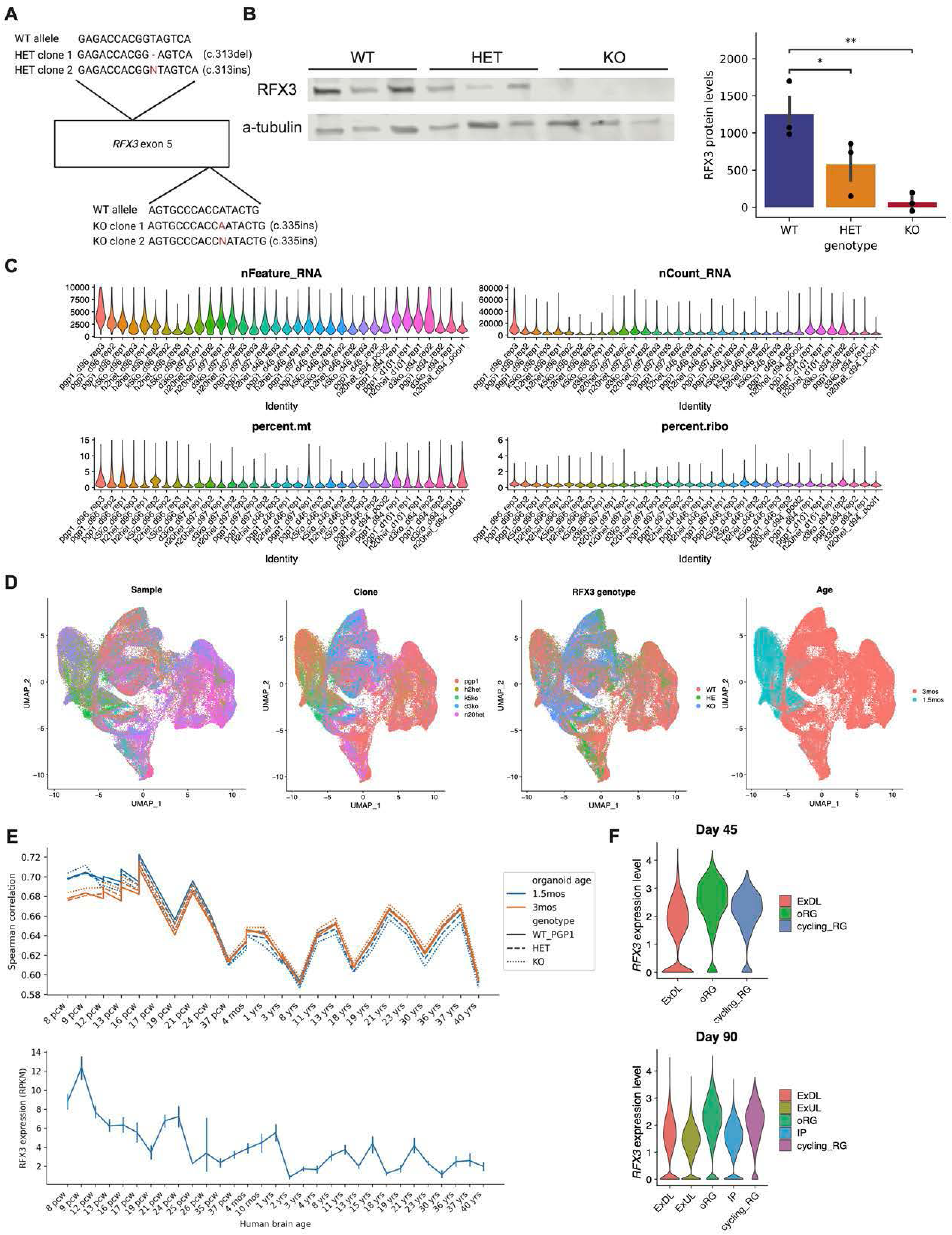
scRNA-seq of *RFX3* WT, HET, and KO dorsal forebrain organoids. A. Generation of *RFX3* isogenic lines (*RFX3* WT, HET, and KO) using CRISPR-Cas9. The schematic shows the indels generated in *RFX3* exon 5. n=2 heterozygous clones, n=2 homozygous clones. Coding nomenclature is for transcript NM_001282116.2. B. Western blot of RFX3 and alpha-tubulin protein levels in *RFX3* WT, HET, and KO iPSCs. C. Violin plot of quality control metrics per sample post-filtering. D. UMAP visualization of meta data. E. Organoid transcriptomes correlate with mid-gestation fetal brain. Spearman correlation coefficient of organoid bulk transcriptome and human brain transcriptome across development (primary motor cortex, BrainSpan) (top). *RFX3* expression in RPKM across human brain development (primary motor cortex, BrainSpan) (bottom). F. *RFX3* expression across cell types in day 45 and day 90 organoids.

**Figure S2:**
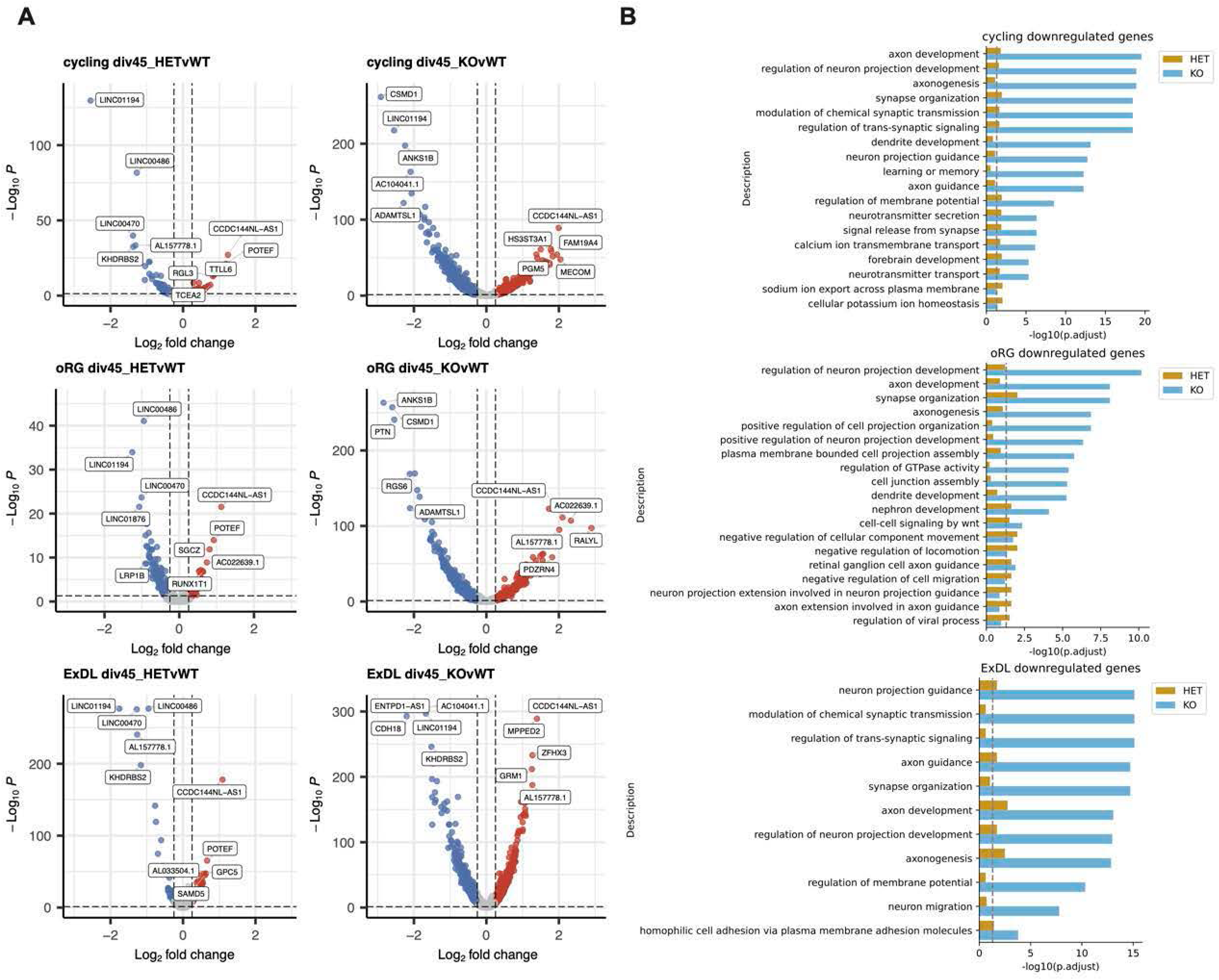
Cell type specific gene expression dysregulation in day 45 *RFX3* deficient dorsal forebrain organoids. A. Volcano plot showing cell type specific differentially expressed genes (DEGs) between *RFX3* HET and WT organoids and *RFX3* KO and WT organoids. Genes significantly downregulated are in blue (log2FC<-0.25, FDR<0.05). Genes significantly upregulated are in red (log2FC>0.25, FDR<0.05). The top DEGs ranked by significance are labeled. B. Gene Ontology (GO) term enrichment analysis of significantly downregulated genes in *RFX3* HET and KO organoids. Benjamini-Hochberg (BH) adjusted p-values calculated with one-sided Fisher’s exact test. Dashed line indicates significance threshold, BH-adjusted p-value=0.05.

**Figure S3:**
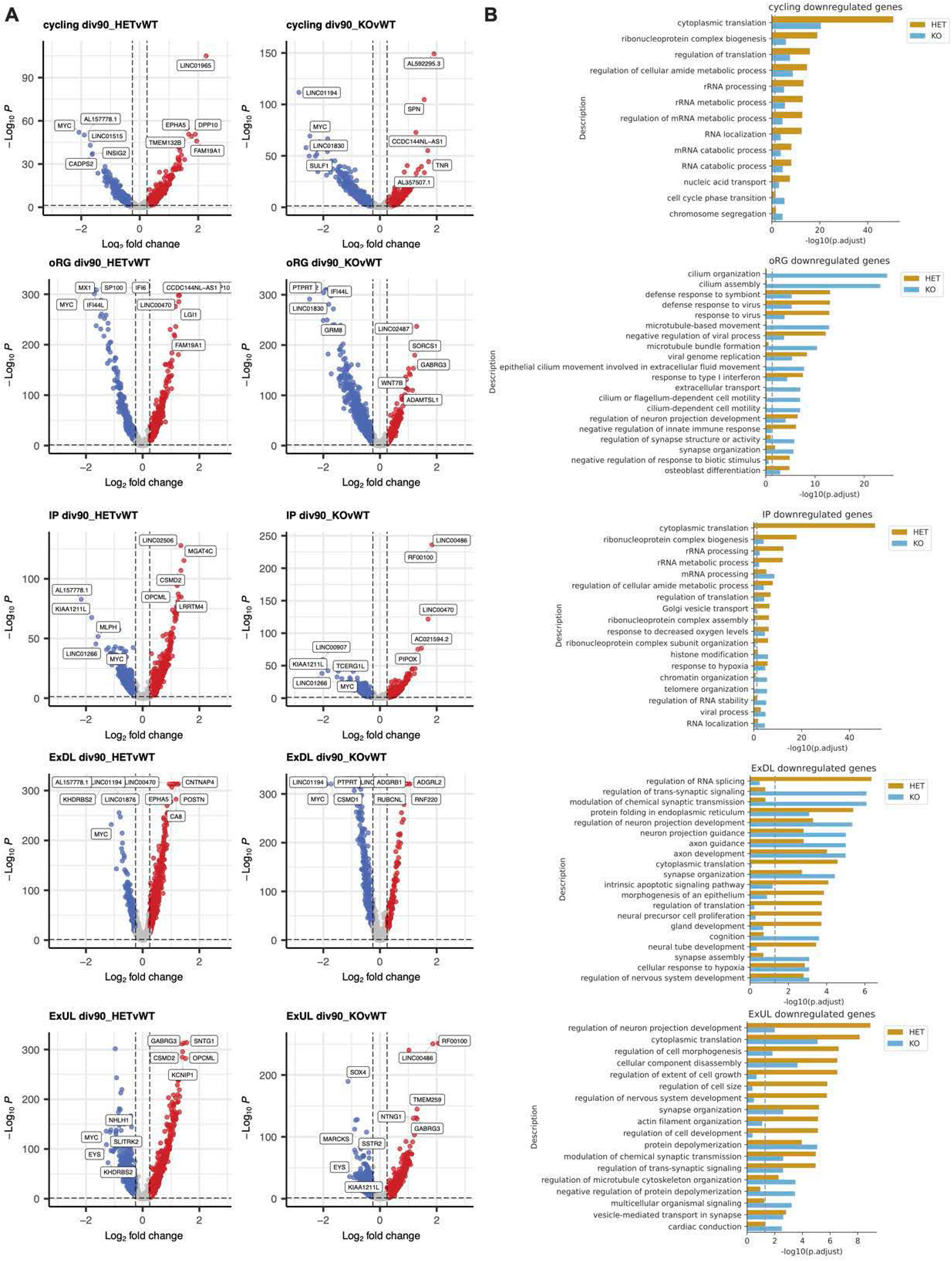
Cell type specific gene expression dysregulation in day 90 *RFX3* deficient dorsal forebrain organoids. A. Volcano plot showing cell type specific differentially expressed genes (DEGs) between *RFX3* HET and WT organoids and *RFX3* KO and WT organoids. Genes significantly downregulated are in blue (log2FC<-0.25, FDR<0.05). Genes significantly upregulated are in red (log2FC>0.25, FDR<0.05). The top DEGs ranked by significance are labeled. B. Gene Ontology (GO) term enrichment analysis of significantly downregulated genes in *RFX3* HET and KO organoids. Benjamini-Hochberg (BH) adjusted p-values calculated with one-sided Fisher’s exact test. Dashed line indicates significance threshold, BH-adjusted p-value=0.05.

**Figure S4:**
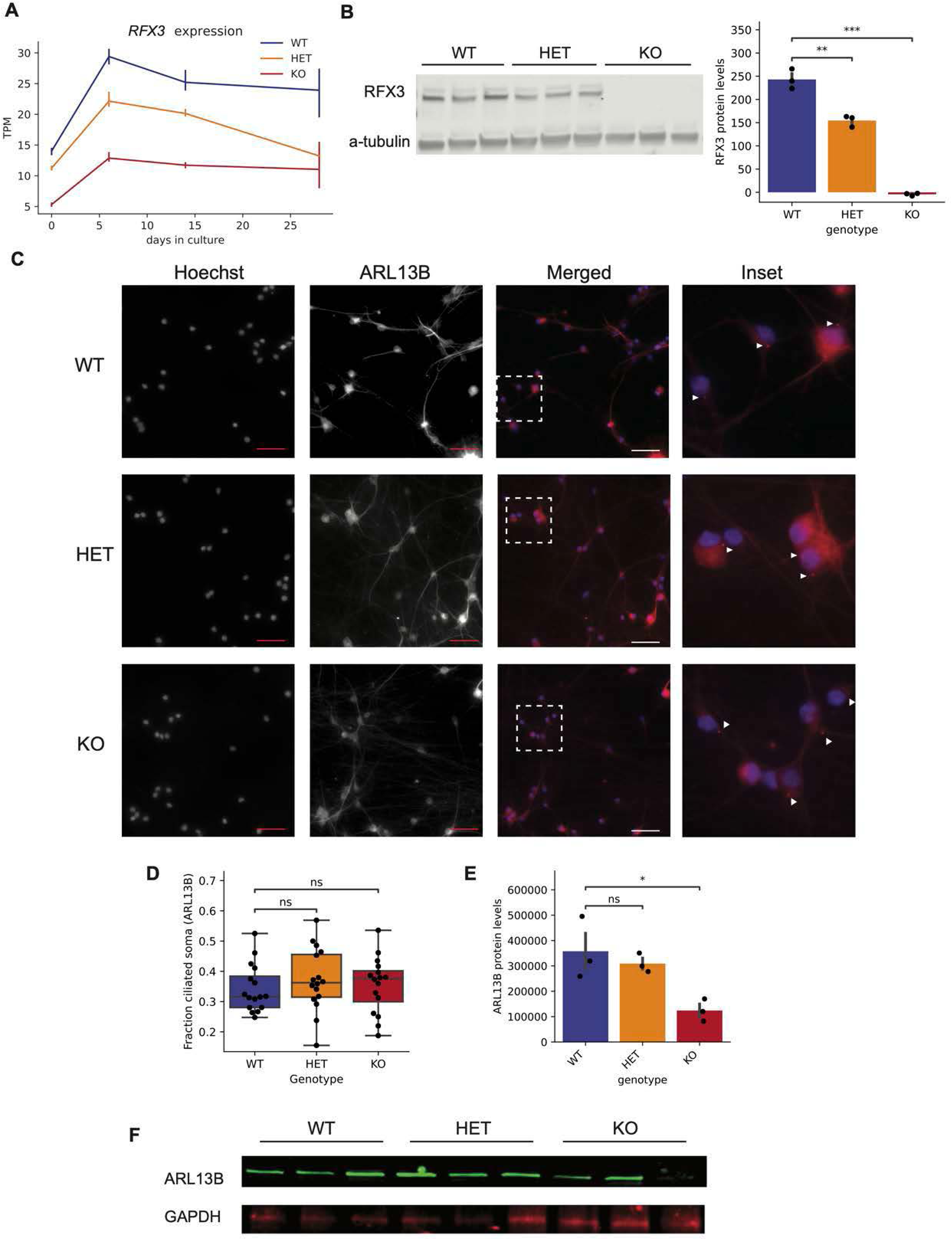
Modeling *RFX3* deficiency in *Ngn2* neurons. A. *RFX3* expression across *Ngn2* iPSC-derived neuron differentiation days 0, 6, 14, and 28 measured by RNA-sequencing. TPM, transcripts per million reads. B. Representative Western blot of RFX3 in day 14 *Ngn2* neurons (left). Quantification of Western blot (right). n=3 independent differentiations per condition. **p< 0.01, ***p<0.005. one-way t-test with Bonferroni correction. C. Representative images of day 20 *Ngn2* neurons stained with Hoechst (nuclei), or ARL13B (cilia). White arrows indicate ARL13B+ cilia. White dashed boxes indicate the inset region. Scale bar=100 um. D. Quantification of fraction of soma with ARL13B+ cilia in day 20 *Ngn2* neurons. n=16 independent wells per condition. ns, not significant, one-way t-test with Bonferroni correction. E. Quantification of ARL13B Western blot. n=3 independent neuron differentiations per condition. Data represent mean +/− SEM. *p <0.05, ns not significant, one-way t-test with Bonferroni correction. F. Representative Western blot of ARL13B and GAPDH in day 14 *Ngn2* neurons. n=3 independent differentiations per condition.

**Figure S5:**
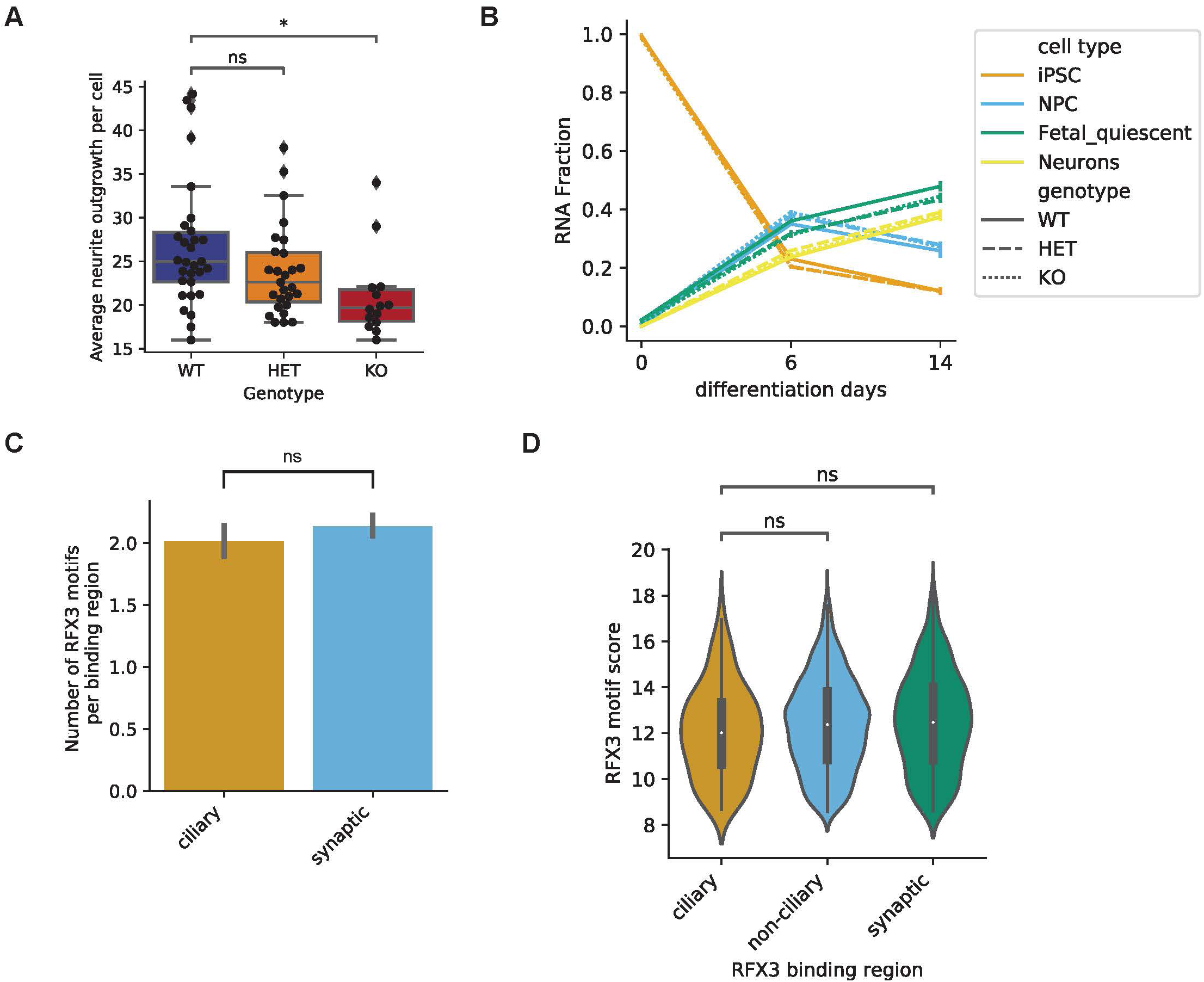
Characteristics of *RFX3* binding motifs in *RFX3* binding sites. A. Average neurite length (um) per cell marked by TUJ1 in day 14 cultures. p <0.05, ns not significant, one-way t-test with Bonferroni correction. B. RNA fraction of cell types estimated by deconvolution of bulk RNA-seq data from iPSC-derived *Ngn2* neurons at days 0, 6, and 14. C. Average number of RFX3 motifs per binding region associated with ciliary genes or synaptic genes. ns, not significant, t-test. Data represent mean +/− SEM. D. Average RFX3 motif score in binding regions associated with ciliary genes, non-ciliary genes, or synaptic genes. ns, not significant, t-test.

**Figure S6:**
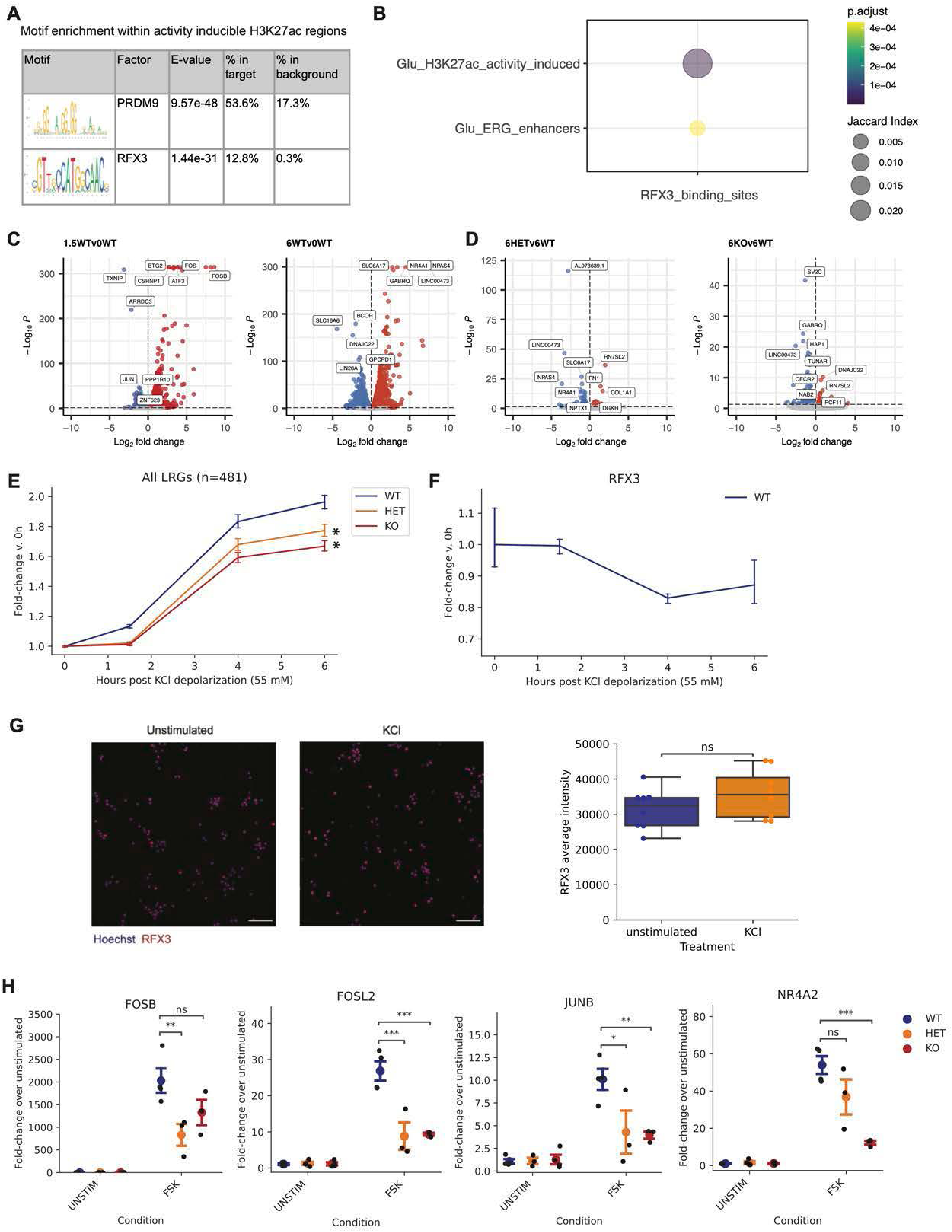
RFX3 modulates CREB-dependent activity-dependent transcriptional responses. A. Top two binding motifs enriched within activity inducible H3K27ac regions in *Ngn2* neurons. B. Jaccard index of RFX3 binding sites and all activity inducible H3K27ac regions or those associated with early response genes (ERG) in glutamatergic (Glu) *Ngn2* neurons. Dot size indicates Jaccard index. Color indicates significance of overlap, two-sided Fisher’s exact test, FDR-adjusted p-value. C. Volcano plot of differentially expressed genes in 1.5 hour or 6 hour KCl stimulated WT vs. unstimulated WT neurons at day 14 in culture. D. Volcano plot of differentially expressed genes in 6 hour KCl stimulated HET vs. WT or KO vs. WT neurons at day 14 in culture. Genes significantly downregulated are in blue (FDR<0.05). (C-D) Genes significantly upregulated are in red (FDR<0.05). Top 10 DEGs are labeled. E. Fold-induction of all late response genes at 1.5, 4, and 6 hours post KCl depolarization compared to unstimulated neurons. Data represented as mean and 95% CI, *p<0.05 t-test, n=3 independent neuron differentiations per genotype. F. *RFX3* expression at 1.5, 4, and 6 hours post KCl depolarization compared to unstimulated WT neurons. Data represented as mean and 95% CI, n=3 independent neuron differentiations. G. Representative images and quantification of RFX3 in unstimulated and KCl stimulated (15 minutes) day 14 WT neurons. n=8 independent wells per condition, ns, not significant, t-test. Scale bar=100 um. H. Fold-induction of *FOSB, FOSL2, JUNB, NR4A2* at 1.5 hours post FSK (10 uM) or H89 (10 uM) pretreatment and FSK (10 uM) assessed by RT-qPCR. Data represented as mean +/− SEM, n=3-4 independent wells per condition. *p<0.05, **p<0.01, ***p<0.005, ns not significant, one-way t-test with Bonferroni correction.

**Figure S7:**
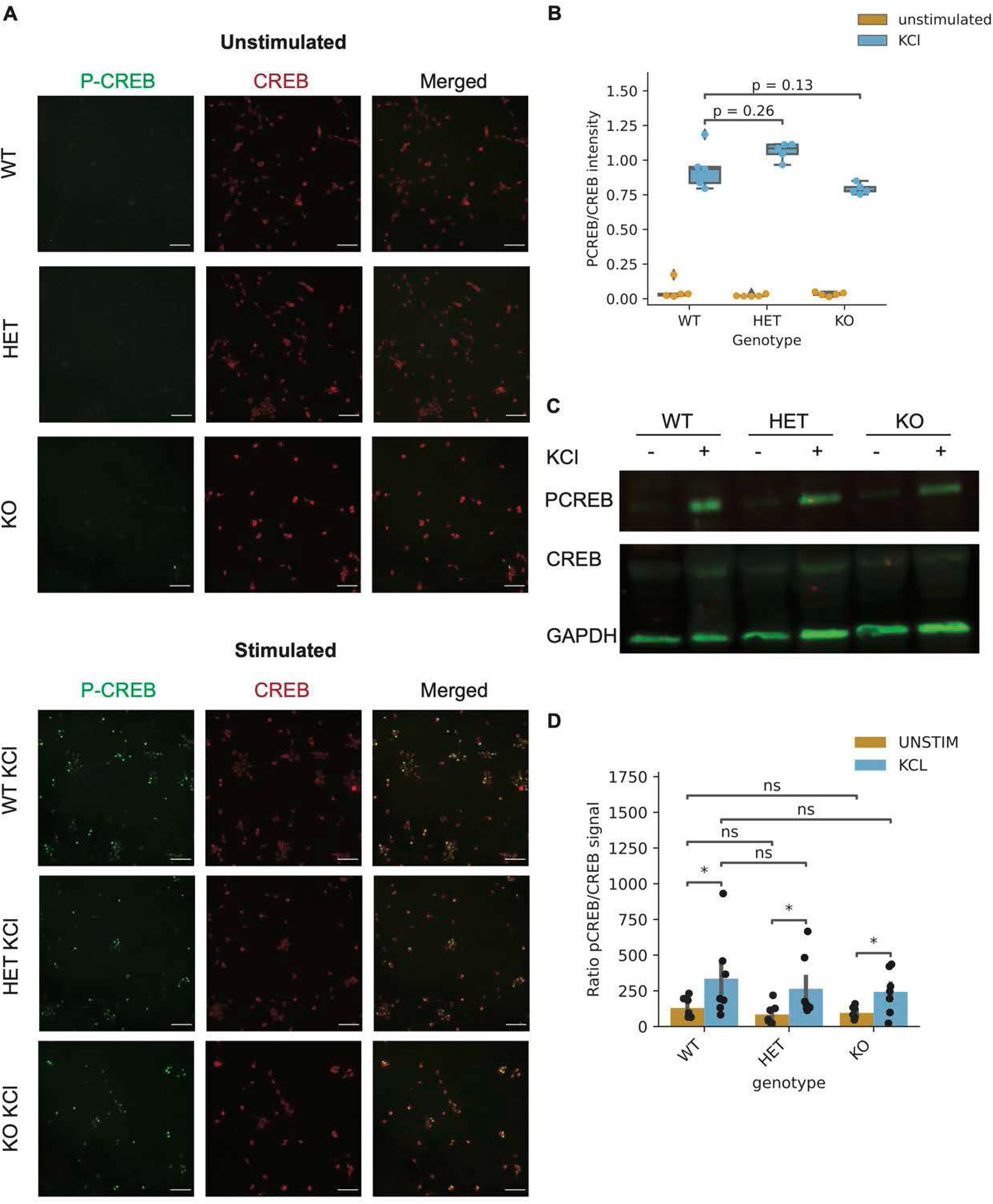
Depolarization induced activation of CREB is intact in *RFX3* deficient neurons. A. Representative images of PCREB and CREB in unstimulated and stimulated (1.5 hours KCl, 55 mM) WT, HET, and KO day 14 neurons. Scale bar=100 um. B. Quantification of nuclear PCREB intensity normalized by CREB intensity in unstimulated and stimulated (1.5 hours KCl, 55 mM) WT, HET, and KO day 14 neurons. P-values correspond to one-way t-test with Bonferroni correction. C. Representative Western blot of PCREB, CREB, and GAPDH in unstimulated and stimulated (15 minutes KCl, 55 mM) WT, HET, and KO day 14 neurons. D. Quantification of Western blot from (C) as the ratio of PCREB/CREB signal. Data represent mean +/− SEM, n=7 replicates per condition. *p<0.05, ns, not significant, one-way t-test with Bonferroni correction.

## Tables

Table S1: Organoid scRNA-seq differentially expressed genes per cell type

Table S2: Organoid scRNA-seq enriched GO terms per cell type

Table S3: RFX3 HET and KO NGN2 neuron differentially expressed genes

Table S4: RFX3 HET NGN2 neuron GO enrichment

Table S5: RFX3 KO NGN2 neuron GO enrichment

Table S6: RFX3 CUT&RUN binding sites

Table S7: RFX3 HET differentially bound sites

Table S8: Immediate early genes (IEGs) and late response genes (LRGs) in WT neurons

Table S9: RFX3 HET and KO neurons differential response to 1.5h KCl depolarization

Table S10: CREB binding sites cobound with RFX3

Table S11: CREB binding sites with diminished CREB binding in *RFX3* HET neurons

## STAR Methods

### Maintenance and culture of iPSCs

Parental PGP1-SV1 iPSC clones and *RFX3* CRISPR-Cas9 edited clones were obtained from Synthego. All iPSC lines were mycoplasma negative, karyotypically normal, and expressed pluripotency markers *OCT4, SOX2, NANOG, SSEA4, TRA-1-60.* iPSCs were maintained in 6-well or 10 cm plates (Corning) coated with LDEV-free hESC-qualified Matrigel (Corning Catalog #354277) in feeder-free conditions with complete mTeSR-plus medium (STEMCELL Technologies) in a humidified incubator (5% CO_2_, 37°C). iPSCs were fed fresh media daily and passaged every 3-4 days.

### Differentiation of iPSCs to neurons

Human iPSC-derived neurons (iNs) were generated from iPSCs transduced with the lentiviruses Tet-O-Ngn2-Puro and FUW-M2rtTA based on a previously published protocol^28^. On day −1 of differentiation, iPSC colonies were dissociated into single-cells using Accutase (STEMCELL Technologies) and 4-8 million cells were plated on Matrigel coated 10 cm dishes with complete mTeSR plus medium supplemented with Y-27632 (10 µM). On day 0, cells were fed N2 media (DMEM/F-12 media, 1X N2, 1X Nonessential Amino Acids) supplemented with doxycycline (2 µg/mL), BDNF (10 ng/mL), NT3 (10 ng/mL), and laminin (0.2 µg/mL). On day 1, media was replaced with N2 media with puromycin (1 µg/mL) in addition to the supplements listed above. On day 2, cells were fed B27 media (Neurobasal media, 1X B27, 1X Glutamax) supplemented with puromycin (1 µg/mL), doxycycline (2 µg/mL), Ara-C (2 µM), BDNF (10 ng/mL), NT3 (10 ng/mL), and laminin (0.2 µg/mL). On day 3, cells were dissociated into single cells using Accutase and replated onto polyethylenimine (PEI)/laminin (0.07% PEI and laminin diluted 1:200 in sterile water) coated 96-well plates (10,000 cells/well) with B27 media supplemented with Y-27632 (10 µM), doxycycline (2 µg/mL), Ara-C (2 µM), BDNF (10 ng/mL), NT3 (10 ng/mL), and laminin (0.2 µg/mL). On days 5, 7, 10, and 14, cells were fed with a half-media change of Conditioned Sudhof Neuronal Growth Medium (1:1 ratio of Astrocyte Conditioned Media and Neurobasal Media, 1X B27, Glutamax, NaHCO3, and transferrin) supplemented with BDNF (10 ng/mL), NT3 (10 ng/mL), and laminin (0.2 µg/mL). Cells were fed with a half-media change of Conditioned Sudhof Neuronal Growth Medium as described above once a week. Cells were maintained in a humidified incubator (5% CO_2_, 37°C).

### Dorsal forebrain organoid culture

Human dorsal forebrain organoids were generated from iPSCs using the STEMdiff Dorsal Forebrain Organoid Differentiation Kit (STEMCELL Technologies Catalog # 08620) according to the manufacturer instructions. On day 0 of organoid formation, iPSCs from each line were dissociated into single cells using Gentle Cell Dissociation Reagent (STEMCELL Technologies Catalog # 100-0485) and 3 x 10^6 cells were seeded into one well of an AggreWell 800 plate (Catalog # 34811) with Seeding Medium. On day 1-5, uniform embryoid bodies were visible and were fed a half-media change of Forebrain Organoid Formation Medium. On day 6, organoids were transferred to 6-well Ultra-Low Adherent plates with Forebrain Organoid Expansion Medium (∼15 organoids per well) and evenly dispersed to ensure no contact between individual organoids and incubated on a stationary level surface. On day 8-25, full media changes with Forebrain Organoid Expansion Medium were performed every 2 days. On day 25, full media changes with Forebrain Organoid Differentiation Medium were performed every 2 days. From day 43 onwards, full media changes with Forebrain Organoid Maintenance Medium were performed every 3-4 days. Organoids were maintained in a humidified incubator (5% CO_2_, 37°C).

### Bulk RNA-sequencing

Total RNA was harvested from samples using the PureLink RNA Mini kit (Life Technologies Catalog # 12183018A). Libraries were prepared with the KAPA mRNA prep. Sequencing was performed on Illumina NovaSeq, 2×150bp configuration at approximately 100X depth per sample. Reads were aligned to hg38 using STAR v2.7.9a^70^ followed by processing with featureCounts (-p -s 2) to obtain a gene-level counts matrix^71^. Differential gene expression analysis was performed using DESeq2^72^ with WT set as the reference condition and results coefficient set as ‘condition_HET_vs_WT’ or ‘condition_KO_vs_WT’, and lfcShrink(type=”apeglm”). Differentially expressed genes with FDR<0.05 and |log2 fold-change| >0.25 were considered significant and used in downstream analysis.

### Isolation of single cells and scRNA-seq library preparation from dorsal forebrain organoids

Individual organoids were dissociated into single cells using 500 µL dissociation solution per organoid (30 units/mL papain, 125 units/mL DNaseI diluted in Hank’s Balanced Salt Solution (HBSS)) for 30 minutes on a shaker in an incubator at 37°C. Organoids were triturated with a P1000 pipette 10-15 times and then returned to the incubator for 15 minutes. Organoids were triturated with a P1000 pipette 10-15 times again to obtain single cells and added to a 15 ml conical tube containing 1 mL of ovomucoid protease inhibitor (10 mg/mL in HBSS). Cells were centrifuged 300g x 5 minutes at room temperature and resuspended in 1 mL HBSS. Cells were counted and assessed for morphology, doublets, and debris. Cell fixation was performed using the Parse Cell Fixation Kit v1 according to manufacturer instructions. Barcoding and library preparation were performed using the Parse Single Cell Whole Transcriptome Kit v1 or Parse Evercode Whole Transcriptome Mega Kit v1 according to manufacturer instructions at the Harvard Single Cell Core. Libraries were paired-end sequenced using Illumina NovaSeq 6000 S2 with target 40,000 reads per cell.

### scRNA-seq analysis

scRNA-seq data for each individual organoid was preprocessed and aligned to the human reference genome Homo_sapiens.GRCh38.93 using the Parse Biosciences computational pipeline split-pipe –mode all. The resulting count matrices were used to create Seurat objects per organoid using Seurat v4^73^. Further quality control removed cells with less than 200 genes per cell, greater than 10000 genes per cell, and greater than 15% mitochondrial content.

SCTransform was used to perform normalization of expression values per sample^74^. After normalization, samples were integrated to align common clusters across individual datasets using Seurat’s integration method^75^. Dimensionality reduction was performed using RunPCA and RunUMAP. A shared nearest neighbor graph was constructed using FindNeighbors with dims 1:30, k.param 100, and cosine metric. Clusters were then identified using the Leiden algorithm at a resolution of 1.5.

Marker genes for each cluster were obtained using FindAllMarkers with a minimum log2 fold-change of 0.25. Cell type identities were assigned to clusters based on reference-based annotations and validation with cell type markers from literature. SingleR^76^ was used for reference-based annotation using reference datasets from the human developing cortex and dorsal forebrain organoids^3,19–21^.

### Differential gene expression and Gene Ontology analysis

Differential gene expression analysis was performed on each cell type using edgeR^77^. The integrated Seurat object was subset to obtain Seurat objects for each cell type. The read counts were modeled and normalized using a negative binomial distribution with the trimmed mean of M-values (TMM) normalization method. To ensure robust signals, only genes that were expressed in at least 5% of one cell type were included in the analysis. The design matrix formula was ∼ genotype + cellular detection rate. Differentially expressed genes between *RFX3* HET or KO and control were identified using the likelihood ratio test (glmLRT). Genes with FDR<0.05 and a log2 fold-change greater than 0.25 were used for downstream analysis.

Gene Ontology (GO) enrichment analysis was performed on the significantly downregulated differentially expressed genes in each cell type using clusterProfiler^78^. GO biological processes with FDR<0.05 were considered significant.

### CUT&RUN-sequencing

Neurons were dissociated into single cells with DNase/papain dissociation solution prepared by combining 20 units/mL papain (Worthington) and 1 unit/mL DNase (Worthington) in DMEM/F12 (STEMCELL Technologies) and incubated at 37°C for 15 minutes. 500,000 single cells per sample were used as input for the CUTANA ChIC/CUT&RUN kit (EpiCypher) according to the manufacturer instructions. Cells were washed twice with CUT&RUN wash buffer and then bound to concanavalin A-coated (conA) beads for 10 minutes at room temperature. ConA bead-bound cells were resuspended in CUT&RUN antibody buffer with the following antibodies: anti-RFX3 (Sigma-Aldrich Cat# HPA035689 diluted 1:100 and Abcam Cat# ab168475 diluted 1:100), anti-H3K4me3 (EpiCypher Cat# 13-0041 diluted 1:100), anti-IgG (EpiCypher Cat# 13-0042 diluted 1:100), anti-CREB (Sigma-Aldrich Cat# 06-863 diluted 1:100) and incubated overnight at 4°C on a shaking nutator with capped ends elevated. After incubation, the conA bead-bound cells were washed twice with Cell Permeabilization Buffer. To each sample, 2.5 µL of pAG-MNase (20X stock) was added followed by incubation for 15 minutes at room temperature. After incubation, the conA bead-bound cells were washed twice with Cell Permeabilization Buffer. On ice, 1 µL of 100 mM Calcium Chloride was added to each sample with gentle vortexing and incubated on a shaking nutator for 2 hours at 4°C. After incubation, 33 µL of Stop Buffer was added to each sample with gentle vortexing. 1 µL (0.5 ng) of *E. coli* Spike-in DNA was added to each sample and incubated for 10 minutes at 37°C in a thermocycler. Samples were placed on a magnetic stand and the supernatant containing CUT&RUN enriched DNA was transferred to clean tubes. DNA was purified from the supernatants using the included DNA Cleanup Columns according to the manufacturer instructions. Purified CUT&RUN-enriched DNA was quantified using Qubit dsDNA HS Assay kit (Thermo Fisher Scientific Cat# Q32851) according to the manufacturer instructions. CUT&RUN-sequencing libraries were prepared using KAPA HyperPrep Kit (Kapa Biosystems Cat# KK8502) according to the manufacturer instructions with the following modifications. 25 µL of purified CUT&RUN-enriched DNA was used as input per sample for End Repair and A-Tailing. To enrich sub-fragments during End Repair and A-Tailing, the thermocycling parameters were 20°C for 30 minutes followed by 50°C for 1 hour. 15 µM adapter stock was used for the Adapter Ligation reaction. Post-ligation cleanup was performed three times with 1X, 1.2X, and 1.2X AMPure beads (Beckman Coulter Cat# A63881). 16 PCR cycles were performed for library amplification. Post-amplification cleanup was performed three times with 1.1X AMPure beads. Purified amplified libraries were quantified by qPCR and library quality and size was assessed using Agilent TapeStation. Libraries were paired-end sequenced with Illumina NextSeq 2×150 bp reads with target 10 million reads per sample. Adapters were trimmed using cutadapt -a AGATCGGAAGAGCACACGTCTGAACTCCAGTCA -A AGATCGGAAGAGCGTCGTGTAGGGAAAGAGTGT (Martin et al., 2011). Alignment to hg38 was performed using bowtie2 --local --very-sensitive-local --no-mixed --no-discordant -- phred33 --dovetail -I 10 -X 700^79^, followed by sorting and indexing with samtools^80^. Peak calling was performed using macs2 callpeak for each sample with -c set as the Rabbit IgG negative control and -f BAMPE -p 0.05^81^. Reproducible peak sets among >3 biological replicates were called using ChIP-R with parameter -m 3^82^. Reproducible peak sets among 2 biological replicates were called using IDR^83^. Peaks were annotated to nearest genes and genomic regions using ChIPseeker^84^. Gene Ontology analysis was performed using clusterProfiler as described above. For categorization of targets as ciliary genes, we used the lists of ciliary genes from CiliaCarta^85^. For categorization of targets as synaptic genes, we used the list of synaptic genes from SynGO^29^. Average peak profiles were generated from RPGC normalized bigwig files using deeptools computeMatrix and plotProfile^86^. Motif enrichment analysis was performed using the MEME suite^87^. Differential motif enrichment analysis was performed using Homer getDifferentialPeaksReplicates.pl^88^.

### Reverse transcription quantitative PCR (RT-qPCR)

RNA was reverse transcribed into cDNA using Cells-to-CT RT buffer and RT enzyme (Thermo Fisher Scientific Cat# A35378) consisting of 22.5 µL RNA and 27.5 µL of Cells-to-CT RT Master Mix. The reverse transcription reactions were incubated at 37°C for 1 hour, and 95°C for 5 min. qPCR was performed using TaqMan Fast Advanced Master Mix (Thermo Fisher Scientific Cat# A35378) and Gene Expression Assays (Thermo Fisher Scientific Cat# 4331182) with 2 µL input cDNA. The thermocycling parameters were 50°C for 2 min, 95C for 2 min, 40 cycles of 95°C for 1 second and 60°C for 20 seconds. TaqMan Gene Expression Assays included *RFX3* Hs01060440_m1, *RFX3* Hs01060430_m1, *FOSB* Hs00171851_M1, *FOSL2* Hs01050117_m1, *JUNB* Hs00357891_s1, *NR4A2* Hs00428691_m1, Human GAPDH Endogenous Control (VIC/MGB probe, primer limited, Thermo Fisher Scientific Cat# 4326317E).

### Western blotting

Cell lysates were collected using RIPA buffer (Boston BioProducts Cat# BP-115D-250) supplemented with Roche cOmplete Protease Inhibitor (Millipore Sigma Cat# 5892791001) and PhosSTOP (Millipore Sigma Cat# 4906845001). Lysates were homogenized with a hand-held spin homogenizer for 10 seconds, then frozen at −80°C for 20 minutes. Lysates were then thawed on ice, centrifuged at 20,000 g for 20 minutes at 4°C, and supernatant was transferred to a clean tube on ice. Lysates were incubated with 4x Laemmli buffer (BioRad), placed in a heat block at 95°C for 5 minutes, then loaded onto 4-15% precast gradient protein gels (BioRad) and separated by electrophoresis (50V for 5 min, 150V for 45 min). Protein samples were then transferred to PVDF membranes and incubated at 4°C overnight with the following primary antibodies: anti-RFX3 (Sigma-Aldrich Cat# HPA035689 diluted 1:100), anti-PCREB (Invitrogen Cat# MA1-114 diluted 1:250), anti-CREB (Sigma-Aldrich Cat# 06-863 diluted 1:500 and Cell Signaling Technology Cat# 4820S diluted 1:500), anti-ARL13B (Proteintech, Cat# 17711-1-AP diluted 1:500), anti-GAPDH (Abcam Cat# ab9485 diluted 1:1,000), anti-alpha Tubulin (Sigma-Aldrich Cat# ab1543 diluted 1:10,000). Membranes were incubated with secondary antibodies diluted 1:5,000 for the target and 1:10,000 for the loading control (GAPDH or alpha Tubulin) and visualized with the Li-Cor Odyssey system and quantified with EmpiriaStudio software (Li-Cor).

### Neuronal depolarization

12 hours prior to depolarization, neuronal cultures were silenced with a full medium replacement consisting of medium supplemented with 1 µM TTX (Abcam Cat# Ab120055) and 100 µM D-APV (Tocris Cat# 0106). Neurons were then left in the silenced condition (unstimulated) or depolarized with KCl depolarization buffer (170 mM KCl, 2 mM CaCl_2_, 1 mM MgCl_2_, 10 mM HEPES, solution pH 7.4) to a final KCl concentration of 55 mM. For pretreatment with H89 dihydrochloride (Tocris Cat# 2910), H89 was added to a final concentration of 10 µM for 1 hour prior to KCl depolarization. For cultures treated with Forskolin (FSK), FSK was added to a final concentration of 10 µM.

### Multielectrode array

48-well CytoView MEA plates (Axion BioSystems) were coated with 0.07% PEI coating buffer at 37C overnight, washed three times with sterile water, air dried, and then coated with Laminin (diluted 1:200 in sterile water) at 37°C overnight. On day 3 of neuron differentiation, wells were plated with 100,000 neurons and 20,000 human iPSC-derived astrocytes (Ncardia Ncyte Astrocytes) per well. Spontaneous neural activity and viability was recorded using the Axion Maestro Pro MEA (Axion BioSystems) with default burst detection settings twice a week for 10 minutes per recording for the following 8 weeks. Half-media changes were performed once a week on a non-recording day with unconditioned neuronal growth medium (consisting of Neurobasal media, B27, NaHCO_3_, Glutamax, Transferrin). Neural activity recordings were processed and analyzed using Axion Axis Navigator and AxIS Metric Plotting Tool softwares with default settings (Axion BioSystems). Data from wells with >0 active electrodes per well were included and were normalized to the number of covered electrodes per well. The number of biological replicates were n=32 wells per genotype from two independent MEA plates.

### Immunostaining

Cells were fixed with 4% paraformaldehyde for 20 minutes at room temperature and stored at 4°C until used for immunostaining. Cells to be stained for PSD95 were fixed with ice cold methanol at −20°C for 20 minutes. Cells were incubated with CellO-IF (Cellorama Tech) for 10 minutes at 37°C, then incubated with primary antibodies diluted in CellO-IF for 1 hour at 37°C. Following incubation, cells were washed with PBS two times, then incubated with secondary antibodies diluted in CellO-IF for 1 hour at 37°C in the dark. Cells were incubated with Hoechst diluted 1:1,000 in PBS for 10 minutes in the dark. After two PBS washes, cells were mounted with glycerol solution (1:1 glycerol and PBS). Primary antibodies included anti-RFX3 (Sigma-Aldrich Cat# HPA035689, diluted 1:100), anti-phospho-CREB (Invitrogen Cat# MA1-114, diluted 1:250), anti-CREB (Sigma-Aldrich Cat# 06-863, diluted 1:500), anti-PSD95 (Neuromab Cat# 75-028, diluted 1:500), anti-SYN1 (Sigma-Aldrich Cat# ab1543, diluted 1:1,000), anti-beta III Tubulin (GeneTex Cat# GTX85469, diluted 1:1,000), anti-ARL13B (Proteintech Cat# 17711-1-AP, diluted 1:800), anti-MAP2 (Abcam Cat# ab5392, diluted 1:2,000). Alexa fluor-conjugated secondary antibodies were used accordingly (diluted 1:500).

### High-content imaging and analyses

96-well plates (Greiner Bio-One Cat# 655090) were imaged with ImageXpress MicroXLS Widefield High-Content Molecular Device microscope (Molecular Device, LLC, MetaXpress v6.6.2.46) under 20X or 40X magnification with 9 evenly spaced fields per well. For Z-stack, five steps and 3 um thickness/step were used, and stacked images were combined with maximum resolution projection. Molecular Devices MetaXpress software was used to design algorithms to systematically identify and quantify selected features from each image. ImageJ software was used to change the LUT for representative images with the same settings applied to each image.

### Statistical analyses

All statistical tests were performed in R (v4.0.1) or Python (v3.8) with multiple hypothesis correction using the Benjamini-Hochberg procedure (FDR<0.05) or Bonferroni correction as specified in figure legends. Statistical tests are noted in figure legends, and include Fisher’s exact test, Student’s t-test, permutation test, and Mann-Whitney U-test.

### Data

Human iPSC-derived NGN2 neuron ATAC-seq data were obtained from GEO: GSE196854 from Sanchez-Priego et al., 2022^37^. H3K27ac CUT&RUN data were obtained from GEO: GSE196207 from Sanchez-Priego et al., 2022^37^.

RNA-seq, scRNA-seq, and CUT&RUN data have been deposited at GEO and are publicly available as of the date of publication. This paper does not report original code. Additional information to reanalyze the data in this paper is available from the corresponding authors upon reasonable request.

